# Novel Binding Partners of the Vacuolar Transporter Chaperone (VTC) complex in Acidocalcisomes of *Leishmania tarentolae*

**DOI:** 10.1101/2025.09.23.677757

**Authors:** Paulina Królak, Orquidea Ribeiro, Bernadette Gehl-Väisänen, Mimmu K. Hiltunen, Adrian Goldman, Keni Vidilaseris

## Abstract

Acidocalcisomes are evolutionarily conserved acidic organelles that are rich in cations and inorganic phosphate, primarily polyphosphates. In kinetoplastid parasites, acidocalcisomes and their polyphosphate content are essential for osmoregulation and environmental adaptation during host switching. In this organelle, polyphosphate is synthesised and transported to the lumen by the vacuolar transporter chaperone (VTC) complex. Interestingly, unlike yeast VTC, which has five components, only two have been observed in kinetoplastids: Vtc1, which contains only a transmembrane domain and Vtc4, which, in addition to a transmembrane domain, also consists of SPX and catalytic domains. In this study, we used proximity-dependent biotinylation (BioID) in *Leishmania tarentolae* to identify proteins located close to the VTC complex. The complex was found near several known acidocalcisomal proteins, including membrane-bound pyrophosphatase (mPPase), vacuolar-type H⁺-ATPase (V-H+-ATPase), Ca²⁺-transporting P-type ATPase (Ca2+-ATPase), zinc transporter (ZnT), and palmitoyl acyltransferase 2 (PAT2). Importantly, this approach revealed three novel VTC binding partners (VBPs) that colocalise and interact with the complex in acidocalcisomes, as confirmed by confocal microscopy, pulldown assays, and AlphaFold3 structural predictions. Together, our results expand the acidocalcisome interactome and suggest that the newly identified VBPs may contribute to the structural organisation and regulatory function of the VTC complex in phosphate homeostasis of kinetoplastid parasites.

**Author summary:** Protozoan parasites such as *Leishmania* and *Trypanosoma* cause serious diseases affecting millions of people worldwide. To better understand how these parasites survive environmental changes during transmission between hosts, we studied a specialised organelle called the acidocalcisome, which stores polyphosphates and helps regulate stress responses. In this work, we used the non-pathogenic *Leishmania tarentolae* as a safe and cost-effective model that shares key cellular features with disease-causing species. Using a combination of CRISPR-Cas9 genome editing, proximity-based labelling (BioID), confocal microscopy, pulldown assays and AlphaFold3 structure prediction, we investigated the vacuolar transporter chaperone (VTC) complex, which synthesises and transports polyphosphate into the acidocalcisome lumen. Proximity proteomics identified several known proteins located near the VTC complex, and importantly, led us to discover three novel proteins that interact with it. These findings open new directions for exploring the organisation and regulation of the VTC complex in protozoan parasites. By revealing novel protein interactions, our study contributes to a deeper understanding of parasite biology and may help identify therapeutic targets for treating neglected tropical diseases.

## Introduction

Neglected tropical diseases, caused by *Leishmania* spp. and *Trypanosoma* spp., pose significant health, social, and economic challenges worldwide (1). These diseases affect millions of people annually: each year there are 700,000 to 1 million cases of leishmaniasis, 6–7 millions of Chagas disease, and up to 1,000 cases of African trypanosomiasis (2). While these infections are primarily endemic to tropical regions, global warming is predicted to alter the distribution and seasonality of their insect vectors, expanding transmission to previously unaffected areas and populations (3).

Kinetoplastid parasites have a unique organelle, the acidocalcisome, which is essential for environmental adaptation and regulation of stress response. Acidocalcisomes play a role in the evasion of host immune defences, ensuring parasite survival during host switching and facilitating the transition between insect and mammalian hosts by maintaining pH homeostasis and osmotic balance (4). They also function as reservoirs of cations and inorganic phosphate (Pi), and take part in calcium signalling (5–7). In *Trypanosoma brucei*, acidocalcisomes are also involved in autophagy (8) and contribute to macromolecular degradation *via* autophagosomes (9). Pi, essential for cellular metabolism, is stored primarily as polyphosphate (polyP), a linear anionic polymer composed of three to thousands of Pi residues linked by high-energy phosphoanhydride bonds (7). PolyP plays a crucial role in osmoregulation and supports parasite survival during changes in environmental conditions associated with host transitions (10, 11). In kinetoplastids, polyP is synthesised by the vacuolar transporter chaperone (VTC) complex, which simultaneously translocates it into acidocalcisomes to prevent cytosolic accumulation, as excess polyP can be toxic (12).

In *Saccharomyces cerevisiae*, the VTC complex consists of five subunits (Vtc1–Vtc5) (11–13). These subunits share a common structural feature: three transmembrane helices (TMHs) at the C-terminus, which are essential for substrate translocation. However, they differ in their additional domains. For example, Vtc1 only contains TMHs, whereas Vtc4 contains, in addition to TMHs, a cytoplasmic catalytic tunnel domain responsible for polyP synthesis and an N-terminal SYG1/Pho81/XPR1 (SPX) domain involved in phosphate sensing and regulation (11–14). Recent cryo-electron microscopy (cryo-EM) studies have shown that the yeast VTC structure is a heteropentameric complex composed of three Vtc1, one Vtc3, and one Vtc4 subunit (15, 16). By contrast, only two VTC subunits, Vtc1 and Vtc4 (**Fig 1A**), have been identified by sequence similarity in kinetoplastids (17–20), though both are important in parasite survival. In *T. brucei*, Vtc1 knockdown disrupts acidocalcisome morphology, impairs osmoregulation, and causes cytokinesis defects, whereas Vtc4 depletion results in impaired growth, virulence, and infectivity (17, 18). In *Leishmania major*, Vtc4 is not required for promastigote viability and proliferation, but its loss delays lesion formation in mice and reduces survival at elevated temperatures (19).

**Fig 1.**
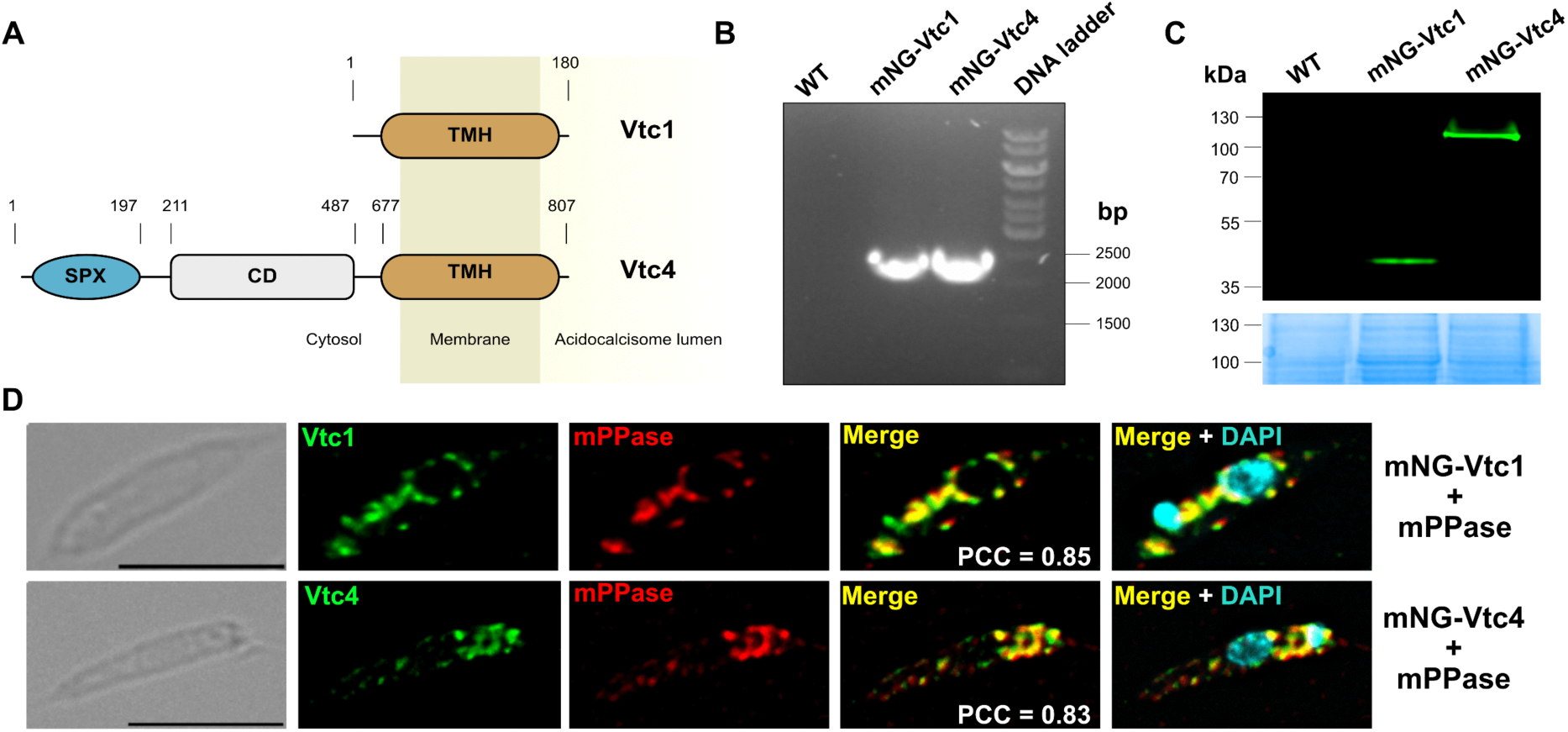
CRISPR/Cas9 tagging and expression of LtVtc1 and LtVtc4. **(A)** Schematic representation of LtVtc1 and LtVtc4. SPX: SYG1/Pho81/XPR domain, CD: Catalytic domain, TMH: Transmembrane helix domain. **(B)** PCR verification of mNG-tagging cassette integration in LtVtc1 and LtVtc4 showing a band at ∼2200 bp, absent in WT. **(C)** Fluorescent detection of mNG-LtVtc1 and LtVtc4 expressed in *L. tarentolae*; Coomassie blue staining of WT *L. tarentolae*, mNG-LtVtc1, and mNG-LtVtc4 confirms comparable total protein levels, ensuring consistent sample loading for fluorescent detection. **(D)** Fluorescence microscopy micrographs of mNG-LtVtc1 and mNG-LtVtc4 with mPPase in *L. tarentolae* (Pearson’s r = 0.85 and 0.83, respectively), scale bar: 5 µm.

Since kinetoplastids have only two VTC components, we speculated that additional, yet unidentified, subunits might contribute to the complex composition and function. Therefore, we used proximity-dependent biotinylation (BioID) (21, 22) to map their protein interaction networks.

Our results demonstrate that the VTC complex is closely associated with several acidocalcisomal proteins, including membrane-bound pyrophosphatase (mPPase), vacuolar-type H⁺-ATPase (V-H+-ATPase), Ca²⁺-transporting P-type ATPase (Ca2+-ATPase), zinc transporter (ZnT), and palmitoyl acyltransferase 2 (PAT2), as well as three proteins of unknown function, which we named ‘VTC Binding Partners’ (VBPs). Immunofluorescence microscopy confirmed the acidocalcisomal localisation of these proteins, encompassing both previously characterised acidocalcisomal components and the newly identified VBPs. Additionally, pulldown assays revealed that VBPs interact with the VTC complex, which is further supported by AlphaFold3 structural predictions.

Together, the identification of novel VTC binding partners lays the groundwork for investigating their contributions to polyphosphate metabolism and acidocalcisome homeostasis in *L. tarentolae*.

## Materials and methods

### L. tarentolae culture

*L. tarentolae* P10 strain promastigotes (LEXSY host P10, Jena Biosciences) were cultured at 27 °C in ventilated tissue culture flasks in 10 ml of brain heart infusion (BHI) medium containing 37 mg/mL BHI powder (Lexsy Broth BHI, Jena Biosciences) supplemented with 5 μg/mL hemin as described by the manufacturer (Jena Biosciences). Cultures were diluted in 1:10 to 1:20 with fresh medium to maintain growth in the mid-log phase at 5 x 10^7^ cells/ml.

### Generation of *L. tarentolae* CRISPR-Cas9 cell lines

An *L. tarentolae* cell line transiently expressing Cas9 and T7 RNA polymerase was generated using the method reported previously (23, 24). Briefly, *L. tarentolae* P10 in the mid-log phase were placed in BHI media and mixed with plasmid pT007 (∼2-5 µg), which encodes for the humanised *Streptococcus pyogenes* Cas9 nuclease gene and T7 RNA polymerase, in a pre-chilled 4 mm electroporation cuvette (BioRad) on ice. Electroporation was performed using the BTX electroporator ECM630 at 1500 V, 25 μF with a double pulse (10 s intervals) and the cuvette was kept on ice for 10 min before being transferred to 10 mL LEXSY Broth BHI supplemented with 5 μg/mL hemin and 50 μg/mL penicillin-streptomycin (Pen-Strep, Jena Bioscience) and incubated at 27 °C. After 24h incubation, 20 μg/mL hygromycin B was added to the culture for polyclonal selection.

### Generation of chromosomally tagged and knockout *L. tarentolae* cell lines

Tagged strains were generated according to the CRISPR-Cas9 method developed by Beneke, Madden (23). Constructs for chromosomal tagging of LtVtc1 and LtVtc4 were amplified by PCR using the pPLOT-mNG-Blast, pPLOT-mCherry-Puro, and pPLOT Puro BirA* plasmids, while gene knockouts were performed using the pTpuro plasmid (23). The generic primer pairs used for LtVtc4 and LtVtc1 were obtained from the LeishGEdit database (www.leishgedit.net, **S1 Table**). These primers are designed to include ∼30 bp homology flanks (HFs), which match sequences flanking the target gene locus. As a result, the PCR product contains a fluorescent tag (with a Myc epitope), an antibiotic resistance cassette, and the necessary homology arms to guide recombination at the intended genomic site. PCR amplification was performed using a two-step protocol. The sgRNA was expressed in vivo from a double-stranded DNA (dsDNA) template containing a T7 promoter, the target-specific sgRNA sequence, and a scaffold region. For transfection, this dsDNA template was delivered alongside the CRISPR repair cassette. N-terminal tagging required a 5′ sgRNA; C-terminal tagging required a 3′ sgRNA; and gene knockout required both sgRNAs and a matching repair cassette containing a puromycin resistance marker (pTPuro plasmid). The sgRNA templates for this CRISPR system were synthesised by PCR from a forward primer that consisted of a T7 promoter sequence followed by the gene-specific target sequence and an overlapping region with the sgRNA scaffold. A generic reverse primer with a 20 bp overlap to the forward primer was then provided, which encoded for the remainder of the scaffold. These short PCR products (ca. 120 bp) were also not purified for transfection into *L. tarentolae* cells.

### Transfection and selection of *L. tarentolae*

*L. tarentolae* cells were transfected as described in the LEXSY protocol (Jena Bioscience). Briefly, the *L. tarentolae* P10 strain was cultured with hygromycin selection over several passages in LEXSY BHI medium and diluted at 1:10 dilution one day before transfection with the pT007 plasmid. We used the BTX electroporator ECM630 at 1500 V, 25 μF with a double pulse in 10 s intervals to transfect the cells with 1 - 10 μg tagging or knockout cassette and the corresponding sgRNA. The cells were then incubated for 20 h in fresh media at 27 °C. Next, 32 μg/mL hygromycin B (Invitrogen, cat 10687010) and 50 μg/mL penicillin-streptomycin (Pen-Strep, Jena Bioscience) and blasticidin (10 μg/mL, Gibco cat 46-1120) or puromycin (10 μg/mL, Gibco cat A11130-03) were added and the culture was continued until the resistant cells emerged. Successful transfection was confirmed by PCR (primers listed in **S1 Table**) and by sequencing (Eurofins, GE).

### Acidocalcisome isolation/preparation

*L. tarentolae* BirA*-LtVtc1 and BirA*-LtVtc4 cell lines, as well as control cells lacking BirA*, were grown in 500 ml of BHI medium supplemented with 50 μM biotin (Sigma-Aldrich, B4501). Biotin was added 24 h after inoculation, and cells were harvested 30 h later.

Cells were collected by centrifugation at 2,000 × g for 3 min, and the pellet (∼2 g wet weight) was washed with cold isolation buffer (125 mM sucrose, 50 mM KCl, 4 mM MgCl₂, 0.5 mM EDTA, 5 mM DTT, 20 mM Hepes-KOH, pH 7.2) supplemented with Pierce Protease Inhibitor Mini Tablets, EDTA-free (Thermo Scientific, A32955). Lysis was performed by grinding for 60 s with silicon carbide (stored at –20 °C before use). The mixture was resuspended in isolation buffer and centrifuged sequentially at 100 × g for 5 min, 300 × g for 10 min, and 1,200 × g for 10 min at 4 °C) to remove silicon carbide and cell debris. The clarified lysate was then centrifuged at 15,000 x g for 10 min at 4 °C to pellet the organelles. The pellet was resuspended in isolation buffer and applied to the 34% step of the discontinuous density gradients containing 20, 24, 28, 34, 37, and 40% (w/V) iodixanol in isolation buffer. Following the published protocol in (6) with one modification (**Panel D in S1 Fig**), we performed single gradient ultracentrifugation instead of two in order to recover not only the acidocalcisome-enriched fraction but also biotinylated proteins from other cellular compartments. The gradient was centrifuged at 50,000 × g for 1 h at 4 °C in an SW Ti rotor. Fractions were collected from the top of the gradient based on their colour and analysed by SDS–PAGE followed by western blotting with anti-mPPase to identify the acidocalcisome-containing fractions (see detailed method below). The rabbit polyclonal anti-TmPPase antibody was generated against purified *Thermotoga maritima* TmPPase (25).

### Western blot analysis

The cells (washed twice in 1× PBS) or the acidocalcisome fractions were lysed on ice for 30 min in RIPA buffer (150 mM NaCl, 20 mM Tris/HCl, pH 7.5, 1 mM EDTA, 1% SDS, and 0.1% Triton X-100) containing a protease inhibitor tablet (Thermo Scientific A32955). Protein concentration was determined *via* Bradford assay (M172-L, VWR Life Sciences). Cell lysates were mixed with 4× Laemmli sample buffer for separation by SDS-PAGE. After SDS-PAGE, the gels were transferred onto nitrocellulose membranes, blocked with 3% bovine serum albumin (BSA) in 1× TBS buffer with 0.5% Tween for 1 h, RT. The blots were incubated with primary rabbit antibodies against TmPPase (1:10,000) (25) and secondary antibodies against anti-rabbit IgG antibody (1:10,000) (sc-2004 by Santa Cruz Biotechnology, INC) or HRP-conjugated avidin (BioRad cat 170-6628) for 1 h. After washing three times with 1× TBS-T, the immunoblots were visualised using Pierce ECL Western blotting substrate according to the manufacturer’s protocol.

### Affinity capture of biotinylated proteins and mass spectrometry

BioID pulldown was done as previously described (21). The acidocalcisome fraction (BirA*-LtVtc1 or BirA*-LtVtc4) was resuspended in 1 mL of lysis buffer (50 mM Tris pH 7.5, 500 mM NaCl, 0.4% SDS, 1 mM DTT). The samples were solubilised in 2% DDM (n-dodecyl-β-D-maltoside) or 2% FC-12 (Fos-choline-12) on ice for 30 min. The samples were centrifuged at 16,000 × g for 10 min at 4 °C, and 100 μL of StrepTactin Sepharose beads (Iba, 2-1201-025) were added to the supernatant and incubated overnight at 4 °C with gentle rotation. The beads were washed three times for 5 min with 1 mL wash buffer (50 mM Tris-HCl pH 7.4, 8 M Urea). The biotinylated proteins bound to the StrepTactin beads were washed eight times with 8 M urea/0.2 M ammonium bicarbonate. Lys-C peptidase was added and incubated o/n at RT in ∼4 M urea/0.1 M ammonium bicarbonate. The peptide digests were centrifuged to pellet the beads, and the supernatant containing the released peptides was collected. Then, trypsin solution was added and the mixture incubated for 4 hrs at 37 °C. The resulting peptide mixtures were desalted using reverse-phase C18 tips (Pierce). Peptides were then vacuum-dried and reconstituted in 1% trifluoroacetic acid (TFA) with brief sonication to ensure they were completely resuspended. Samples were analyzed by liquid chromatography–mass spectrometry (LC-MS). The acquired MS data were searched against TriTrypDB databases (www.tritrypdb.org) (26) for *L. tarentolae* Parrot Tar II 2019 (27), using Thermo Proteome Discoverer 2.5.

The protein list from the mass spectrometry data was screened for overlap and manually checked for homology to *T. brucei* proteins using the TriTrypDB-8.1 TREU 927 database (https://tritrypdb.org/tritrypdb/app). Their potential subcellular localisation was analysed using the TrypTag database (http://www.tryptag.org/) (28), a trypanosome genome-wide protein localisation resource. Hits localised to small cytoplasmic organelles (29) were further examined using published proteomes of the acidocalcisome (6), glycosome (30, 31), and RNA-granule/polysomes (32) from *Leishmania* and *Trypanosoma*. Additional checks were performed to assess potential localisation in the mitochondria (33, 34), nucleus (35), and for polyP binding capacity (36), based on available proteomic datasets. Raw mass spectrometry data were deposited in the PRIDE repository (ProteomeXchange Consortium) under accession number PXD075501.

### Pulldown assays

*L. tarentolae* cultures (10 mL) expressing mNG-tagged LtVtc4 and mCh-tagged binding partner candidates were pelleted by centrifugation (4000 × g, 10 min at 4 °C ) and washed with 1× PBS. The cell pellets were lysed by seven freeze-thaw cycles in 100 µL of lysis buffer (50 mM HEPES pH 7.5, 150 mM NaCl, 1 mM IP_6_ (Inositol hexaphosphate, Sigma-Aldrich), 1mM DTT, 1 mM PMSF, 2 µg/mL Pepstatin A (Sigma Aldrich, P5318-5MG) , and Pierce Protease Inhibitor Mini Tablets, EDTA-free (Thermo Scientific A32955)).

Membrane proteins were solubilised by adding 1% DDM, followed by incubation on a roller (1.5 h, 4 °C). After centrifugation (25,000 × g, 1 h at 4 °C ) to remove cell debris, the supernatants were incubated on a roller (o/n, 4 °C) with 25 µL of ChromoTek mNeonGreen-Trap Agarose beads (Proteintech, AB_2827593). Additionally, for mNG-LtVtc4 + mCh-LtVBP3, ChromoTek RFP-Trap Magnetic Particles M-270 (Proteintech, AB_2861253) were used. Beads were collected by centrifugation (1000 × g, 1 min) using a spin column, washed three times with 1 mL of lysis buffer containing 0.05% DDM (centrifugation at 1000 × g, 1 min per wash). For magnetic beads, a magnetic stand was used for collection and washing. Finally, beads were resuspended in 50 µl of lysis buffer containing 0.05% DDM. Samples from each purification step were resolved by SDS-PAGE and analysed by in-gel fluorescence using the Sapphire FL Biomolecular Imager (Azure Biosystems). Fluorescence was detected using 488 nm excitation for mNG and 532 nm for mCh, which also partially excites the mNG emission signal due to spectral overlap. As a result, some mNG signals may appear yellowish in the 532 nm channel (see **S2 Fig**). A 638 nm channel was used to detect the protein marker (PageRuler Plus Prestained Protein Ladder, 26619, ThermoFisher Scientific).

### Membrane preparation and purification of the VTC complex from *L. tarentolae*

For *in vitro* assessment of polyP polymerase activity, the VTC complex was purified from *L. tarentolae* expressing N-terminally c-Myc–mNG-tagged LtVtc4. Frozen cell pellets (corresponding to 1 L culture) were thawed and resuspended in buffer A (50 mM HEPES pH 7.5, 300 mM NaCl, 0.3 M sucrose, 1 mM MnCl₂, 1 mM MgCl₂), supplemented with 1 mM DTT, 1 mM PMSF, 2 µg/mL Pepstatin A (Sigma Aldrich, P5318-5MG) and Pierce Protease Inhibitor Mini Tablets, EDTA-free (Thermo Scientific A32955), to a final volume of 50 mL. Cells were lysed by high-pressure homogenisation using an Emulsiflex system. Cell debris was removed by centrifugation (4000 rpm, 10 min at 4 °C), and the clarified lysate was transferred to ultracentrifuge tubes. Membrane fractions were pelleted by ultracentrifugation using a Beckman Ti45 rotor (42,000 rpm for 1 h at 4 °C). The membrane pellet was resuspended in buffer A using a Dounce homogeniser and subjected to a second ultracentrifugation step under the same conditions to further enrich membrane components. The resulting membrane pellet was resuspended in buffer B (50 mM HEPES pH 7.5, 300 mM NaCl, 10% glycerol), supplemented with 1 mM DTT, 1 mM PMSF, 2 mM IP₆ (Inositol hexaphosphate, Sigma-Aldrich), 2 µg/mL Pepstatin A (Sigma Aldrich, P5318-5MG), and Pierce Protease Inhibitor Mini Tablets, EDTA-free (Thermo Scientific A32955), using a Dounce homogeniser.

Membrane fractions were adjusted to a final total protein concentration of 20 mg/mL using solubilization buffer (50 mM HEPES pH 7.5, 300 mM NaCl, 10% glycerol), supplemented with 2 mM IP₆ (Inositol hexaphosphate, Sigma-Aldrich), 1 mM DTT, 1 mM PMSF, Pepstatin A (Sigma Aldrich, P5318-5MG), Pierce Protease Inhibitor Mini Tablets, EDTA-free (Thermo Scientific A32955) and solubilised under native conditions with 1% DDM, followed by incubation on a roller (1.5 h, 4°C). Insoluble material was removed by ultracentrifugation using a Beckman Ti50.2 rotor (33,000 rpm, 1 h at 4 °C). The clarified lysate was applied to anti-c-Myc agarose resin (Pierce Anti-c-Myc Agarose, 20168) using a gravity-flow column. Following binding, the resin was washed with solubilisation buffer containing reduced detergent concentration (0.05% DDM) to remove non-specifically bound proteins.

The VTC complex was eluted using competing c-Myc peptide (Pierce c-Myc Peptide, 20170), and elution fractions were concentrated using a 100 kDa MWCO Amicon Ultra centrifugal filter (Millipore, UFC8100). All purification steps were performed at 4 °C. Samples from purification were resolved by SDS-PAGE and analysed by in-gel fluorescence using the Sapphire FL Biomolecular Imager (Azure Biosystems), followed by Coomassie Brilliant Blue staining. Fluorescence was detected using 488 nm excitation for mNG. Gel images were acquired using a Bio-Rad Gel Doc EZ Imager. Purified VTC complex was used directly for subsequent polyP activity assays.

### PolyP activity assay and urea-PAGE analysis

PolyP polymerase activity of purified VTC complex was assessed using an enzymatic assay followed by urea-PAGE analysis. Reactions were performed in a buffer containing 50 mM HEPES pH 7.5, 300 mM NaCl, 2 mM MgCl_2_, 2 mM MnCl_2_, and 0.05% DDM. Purified protein (1mg/mL, 10 µL) was mixed with 30 µL assay buffer, and the reaction initiated by the addition of 10 µL substrate mixture containing 25 mM ATP and 25 mM IP₆ (Inositol hexaphosphate, Sigma-Aldrich). Reactions were incubated at 30 °C for 2 h or overnight. PolyP products were analysed by urea-PAGE. Samples were mixed with DNA loading dye (6X TriTrack DNA loading dye, Thermo Scientific, R1161) and separated on pre-run urea gels in 1× TBE buffer at 150 V. Gels were pre-run for 20 min prior to sample loading and electrophoresed for 40 min. PolyP standard (0.1 µg) (Sodium polyphosphate, EMPLURA, 10361-03) was included for size comparison. Following electrophoresis, gels were stained with DAPI-containing buffer (2 µg/mL DAPI in 20% methanol, 5% glycerol, 50 mM Tris pH 10.5) for 1 h in the dark, followed by destaining in buffer lacking DAPI (20% methanol, 5% glycerol, 50 mM Tris pH 10.5) for 1 h in the dark. PolyP was visualised using Bio-Rad Gel Doc EZ Imager based on DAPI fluorescence shift upon binding.

### Immunofluorescence microscopy and statistics

Cells were attached to coverslips by centrifugation (3,000 × g, 5 s) and fixed with 4% (w/v) paraformaldehyde (20 min, RT). After fixation, cells were washed with 1× PBS, permeabilised with 0.25% Triton X-100 in 1× PBS (V/V) for 5 min at RT, washed again with 1× PBS, and blocked with 3% BSA in 1× PBS (w/v) (30 min, RT). The coverslips were incubated with primary antibodies (ChromoTek mouse anti-mNeonGreen (1:250, 32F6, Proteintech), Rabbit Polyclonal anti-mCh (1:400, 26765-1-AP, Proteintech)) diluted in 1× PBS in a humidified chamber (1 h, RT), washed three times with 1× PBS, and then incubated with secondary antibodies (CoraLite 488-Conjugated Goat Anti-Mouse IgG(H+L) (1:750, SA00013-1, Proteintech), Goat anti-Rabbit IgG (H+L) Cross-Adsorbed Secondary Antibody, Alexa Fluor 594 (1:1000, A-11012, ThermoFisher Scientific)) diluted in 1× PBS (1 h, RT). After three washes with 1× PBS, the coverslips were rinsed with MilliQ water and mounted on glass slides using Fluoromount G + DAPI (Southern Biotech).

Cells were imaged using a Leica Stellaris 8 confocal microscope with a DMI8 base, operated using LAS X 3.5.2 acquisition software. The objective used was an HC PL APO CS2 40×/1.25 GLYC (Glycine Immersion Medium). Fluorescence detection was performed using hybrid detectors (HyD) and solid-state lasers, with the following settings: mNG detected using Alexa 488 (HyD X2; emission range: 510–540 nm), DAPI detected using Alexa 405 (HyD S1; emission range: 420–490 nm), and mCh detected using Alexa 568 (HyD S3; emission range: 590–740 nm). The pinhole size was set to Airy 1, and the scan speed was 400 Hz.

Images were acquired with these settings by capturing Z-stacks ranging from 8 to 18 slices, with slice spacing between 2.16 and 6.13 µm and pixel size ranging from 56.78 to 116.44 nm. Images were processed using FIJI (ImageJ) (37), and deconvolution was performed for 10 cycles using the DeconvolutionLab2 plugin (38). Pearson’s correlation coefficients were calculated using the Coloc2 plugin (37) by analysing Z-stacks. For each condition, 30–50 randomly selected cells were analysed, depending on the experiment. Statistical comparisons of Pearson’s correlation coefficient values were performed in Python using SciPy, statsmodels, and Pingouin packages (39). One-way ANOVA and Tukey’s honestly significant difference (HSD) post hoc test were used for comparisons.

### Gene ontology analysis

GO analysis of biological processes, cell components, and molecular functions was performed on the 412 overlapping LtVtc1 and LtVtc4 interactors using the TriTrypDB GO analysis tool (40) with the complete *L. tarentolae* Parrot Tar II 2019 (ATCC 30267) genome as the reference. Fisher’s exact test was applied, with a *p*-value cutoff of 0.01, and significance was defined as Bonferroni-adjusted *p*-value < 0.01 and Benjamini-Hochberg FDR < 0.01.

### Protein structure prediction and analysis

Protein structure predictions were performed using AlphaFold3 (41) and ColabFold for actifpTM analysis (42, 43). Protein transmembrane topology and orientation were predicted using MEMSAT-SVM *via* the PSIPRED server (https://bioinf.cs.ucl.ac.uk/psipred/) (44, 45). Molecular analysis and molecular graphics were produced in PyMOL (https://pymol.org). Protein sequence similarity searches were performed using BLAST in the UniProt database (46). Sequence alignment was performed using Clustal Omega (47) and displayed using ESPript 3.0 (48).

## Results

### Localisation of *L. tarentolae* Vtc1 and Vtc4 to acidocalcisomes

To investigate the localisation of LtVtc1 and LtVtc4 in *L. tarentolae* (orthologous to Tb927.7.3900 and Tb927.11.12220 in *Trypanosoma brucei*, respectively), we tagged both proteins with mNeonGreen (mNG) using CRISPR-Cas9, following the method of Beneke *et al*. (23), which has been successfully applied in *L. tarentolae* (24). PCR analysis showed clear bands at ∼2200 bp, corresponding to integration of the mNG-tagging cassette into the respective VTC subunit loci, with no amplification in the wild-type (WT) control, confirming successful chromosomal tagging (**Fig 1B**). Expression of mNG-LtVtc1 and mNG-LtVtc4 was confirmed by SDS-PAGE followed by fluorescent imaging (**Fig 1C**). Immunofluorescence microscopy showed that both fusion proteins colocalised with the acidocalcisomal marker mPPase, confirming their acidocalcisomal localisation (**Fig 1D**). To verify that the VTC complex remains functionally active, the complex was purified from *L. tarentolae* and its polyP polymerase activity was assessed (**panel A-B in S3 Fig**). PolyP synthesis was confirmed by urea-PAGE analysis, demonstrating that the natively isolated complex retains enzymatic activity (**panel B in S3 Fig**).

### Identification of proteins in close proximity to the VTC complex

After confirming the acidocalcisomal localisation of LtVtc1 and LtVtc4, both genes were N-terminally tagged with BirA* to enable proximity labelling (BioID) of potential VTC interactors from the cytoplasmic side. BioID is an established technique that identifies proteins within a ∼10 nm radius of the bait protein in live cells, oÅering insights into potential protein-protein interactions (21, 22).

Successful integration of the BirA* cassette into the chromosomes and expression of the fusion proteins were verified by PCR and western blot, respectively (**panel A-B in S1 Fig**). In the presence of biotin, *L. tarentolae* cells expressing BirA*-LtVtc1 and BirA*-Vtc4 showed strong enrichment of biotinylated proteins (**panel C in S1 Fig**). Following cell lysis, iodixanol-gradient ultracentrifugation (**panel D in S1 Fig**) yielded acidocalcisome-enriched fractions (**panel E in S1 Fig**), with BirA*-LtVtc1 and BirA*-LtVtc4 predominantly localising to fractions 4 and 5, respectively (**panel F-G in S1 Fig**). The slight difference in fractionation is due to variations in the collected gradient volumes.

Acidocalcisome fractions from BirA*-LtVtc1 and BirA*-LtVtc4 cell lines were solubilised with 2% DDM or 2% FC-12, based on detergent solubilisation screening of mNG-LtVtc4 showing that these detergents efficiently extracted more than 50% of the LtVtc4 (**panel H in S1 Fig**). Affinity purification of the biotinylated proteins followed by liquid chromatography-mass spectrometry (LC-MS) resulted in a total of 3,160 protein identifications across four datasets, corresponding to 1,240 non-redundant (unique) protein hits (**Fig 2A and S2-S5 Table**). From the BirA*-LtVtc1 samples, 912 proteins were identified in the DDM-solubilised and 752 in the FC-12-solubilised fractions, while BirA*-LtVtc4 samples yielded 929 and 567 proteins, respectively (**Fig 2A**). As expected, there was substantial overlap between the conditions, with 414 proteins detected in all datasets (**Fig 2A**).

**Fig 2.**
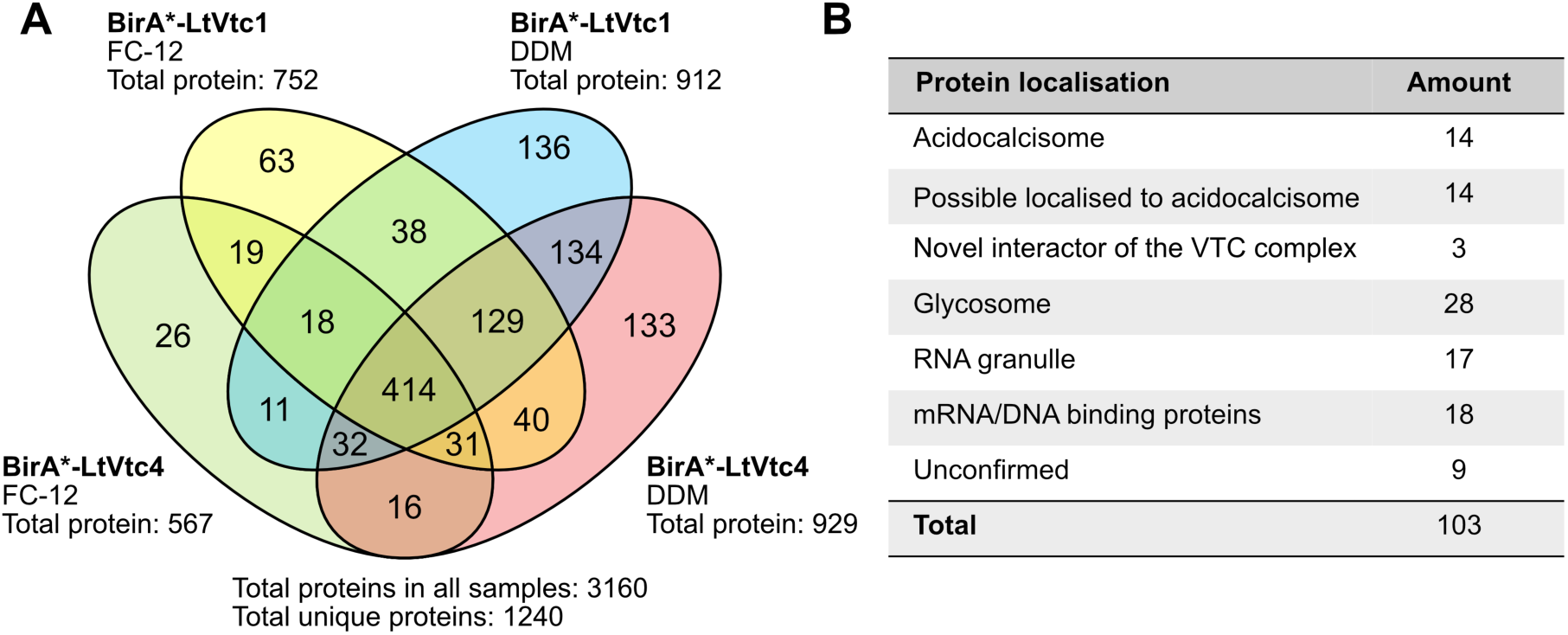
Number of proteins observed in the proximity-dependent biotinylation experiment. **(A)** Venn diagram showing overlap of proteins observed in the four datasets (BirA*-LtVtc1, FC-12; BirA*-LtVtc4, FC-12; BirA*-LtVtc1, DDM; BirA*-LtVtc4, DDM). **(B)** Predicted localisation of *L. tarentolae* BioID hits based on the localisation of their *T. brucei* orthologs in the TrypTag database (http://www.tryptag.org/).

To further characterise the overlapping proteins, we performed gene ontology (GO) enrichment analysis using the *L. tarentolae* Parrot Tar II 2019 genome as reference in the TriTrypDB GO analysis framework (26, 40) to identify enriched biological processes and functions among proteins proximal to the VTC complex. This analysis revealed significant enrichment across multiple GO categories (**S4 Fig**). Among these, ATP metabolic process (GO 0046034), intracellular organelle (GO 0043229) and proton channel activity (GO 0015252) were notably overrepresented in the biological processes, cell components and molecular functions categories, respectively.

In parallel, we assessed the predicted subcellular localisation of the *L. tarentolae* BioID hits by mapping them to their *T. brucei* orthologs annotated in the TrypTag database, a genome-wide protein localisation resource for *T. brucei* (28), together with published organelle proteomes (6, 30–32). Many of the identified *L. tarentolae* proteins were assigned to acidocalcisomes based on this orthology-based analysis, whereas others were associated with glycosomes and RNA granules (**Fig 2B and S6 and S7 Table**). Some of these non-acidocalcisomal assignments may reflect contamination during acidocalcisome isolation. However, the high number of non-acidocalcisomal interactors, together with the known presence of polyP in other organelles (36), prompted further analysis.

PolyP-binding proteins were identified by cross-referencing the *L. tarentolae* BioID datasets with proteins previously reported to interact with polyP in trypanosomatids (36). This analysis identified 50 putative polyP-binding proteins, which were assigned to acidocalcisomes (11 %), glycosomes (12 %), nucleoplasm (12 %), other compartments (20 %), and the cytoplasm (45 %), with ribosomal subunits being highly represented (**S8 Table**). GO enrichment analysis further showed that translation initiation factors and ribosomal proteins were overrepresented among the BioID hits.

In addition to the polyP-binding and translation-related proteins, the BioID datasets also contained proteins assigned to RNA granules based on TrypTag localisation data. Seventeen *L. tarentolae* BioID hits were mapped to *T. brucei* orthologs predicted to localise to RNA granules, which are cytosolic, non-membrane-bound structures involved in RNA storage, processing, and degradation (**S7 Table**). Among these, Alba domain-containing protein 3 (LtALBA3, LtaPh_3424900), Poly(A) binding protein 2 (LtPABP2, LtaPh_2500900) and Zinc finger protein family member (ZC3H41, LtaPh_2713900) were consistently detected across all VTC BioID datasets. LtALBA3 and LtPABP2 also showed high peptide coverage (**S7 Table**), and their orthologs have been reported in the acidocalcisome proteome of *T. brucei* (6).

### Proximal acidocalcisome proteins and novel interactors of the VTC Complex in *Leishmania tarentolae*

To identify relevant protein interactions within acidocalcisomes, we manually filtered the 1240 unique *L. tarentolae* BioID hits by mapping them to their *T. brucei* orthologs and examining their predicted localisation in small cytoplasmic organelles (acidocalcisomes, glycosomes, lipid droplets, and RNA granules) (29) using the TrypTag database (http://www.tryptag.org/) (28). This analysis identified 103 *L. tarentolae* proteins whose *T. brucei* orthologs localised to these organelles (F**ig 2B** **and S9 Table**). Among these, 14 proteins had previously been reported to localise in *T. brucei* acidocalcisomes (7, 49, 50), including eight proteins consistently detected across all BioID datasets: mPPase, Ca^2+^-ATPase, ZnT, PAT2, and two V-H^+^-ATPase subunits (-A, -c putative subunit), as well as Vtc1 and Vtc4.

V-H^+^-ATPase is a multi-subunit proton pump consisting of two sub-complexes, the peripheral V1 complex (composed of eight subunits: A-H), responsible for ATP synthesis/hydrolysis, and the integral-membrane V0 complex (composed of five subunits: a, c, c’, c”, d, and e), responsible for proton translocation (51). In addition to the two V-H^+^-ATPase subunits present in all datasets, subunits D and a occurred in one and three datasets, respectively (**Table 1**). These findings suggest that the VTC and V-H^+^-ATPase complex are close to each other, although not all subunits may be accessible to BirA* biotinylation. Supporting this, VTC1 deletion in yeast decreased the levels of several V- H^+^-ATPase subunits and impaired proton uptake activity, indicating that Vtc1 is important in maintaining V-ATPase stability (52).

**Table 1.** Acidocalcisome proteins near the VTC complex.

| No. | Gene |  | Protein name | Membrane association | Peptide counts (% Coverage) |  |  |  |
| --- | --- | --- | --- | --- | --- | --- | --- | --- |
|  | *L. tarenolae* | *T. brucei* |  |  | DDM |  | FC-12 |  |
|  |  |  |  |  | BirA*-LTVTC1 | BirA*-LTVTC4 | BirA*-LTVTC1 | BirA*-LTVTC4 |
| 1 | LtaPh_1410200 | Tb927.7.3900 | Vacuolar transporter chaperone 1 (Vtc1) | Yes | 5 (34.0) | 1 (8.3) | 2 (14.0) | 1 (8.3) |
| 2 | LtaPh_0902200 | Tb927.11.12220 | Vacuolar transporter chaperone 4 (Vtc4) | Yes | 25 (53.3) | 18 (30.3) | 5 (9.5) | 20 (32.4) |
| 3 | LtaPh_1706300 | Tb927.8.1160, Tb927.8.1180, Tb927.8.1200 | P-type Ca<sup>2+</sup>-ATPase | Yes | 2 (3) | 3 (3) | 2 (3) | 1 (3) |
| 4 | LtaPh_2829500 | Tb927.11.11160 | Phosphate transporter 91 (Pho91) | Yes | 1 (2) | 0 (0) | 0 (0) | 0 (0) |
| 5 | LtaPh_3111800 | Tb927.4.4380 | Membrane bound pyrophosphatase (mPPase) | Yes | 4 (6) | 4 (7) | 5 (7) | 5 (6) |
| 6 | LtaPh_3128400 | Tb927.4.4960, Tb927.8.7460 | Zinc transporter (ZnT) | Yes | 3 (17) | 2 (8) | 1 (3) | 1 (3) |
| 7 | LtaPh_3625400 | Tb927.10.7020 | Acid phosphatase (AP) | No | 0 (0) | 1 (5) | 0 (0) | 0 (0) |
| 8 | LtaPh_2121161 | Tb927.10.200, Tb927.11.7480 | V-type H<sup>+</sup>-ATPase subunit c (V-H<sup>+</sup>-ATPase_c) | Yes | 1 (8) | 1 (8) | 1 (8) | 1 (8) |
| 9 | LtaPh_0512200 | Tb927.5.550 | V-type H<sup>+</sup>-ATPase subunit D (V-H<sup>+</sup>-ATPase_D) | No | 1 (8) | 0 (0) | 0 (0) | 0 (0) |
| 10 | LtaPh_2319200 | Tb927.5.1300 | V-type H<sup>+</sup>-ATPase subunit a (V-H<sup>+</sup>-ATPase_a) | Yes | 2 (6) | 1 (3) | 1 (2) | 0 (0) |
| 11 | LtaPh_3435900 | Tb927.4.1080 | V-type H<sup>+</sup>-ATPase subunit A (V-H<sup>+</sup>-ATPase_A) | No | 1 (3) | 2 (5) | 2 (5) | 1 (2) |
| 12 | LtaPh_1512200 | Tb927.9.5490 | Magnesium transporter (MgT) | Yes | 1 (2) | 2 (4) | 0 (0) | 0 (0) |
| 13 | LtaPh_3302521 | Tb927.10.10800 | Palmitoyl acyltransferase 2 (PAT2) | Yes | 2 (6) | 1 (3) | 1 (3) | 1 (3) |
| 14 | LtaPh_0904900 | Tb927.11.12490 | Inward-rectifier K<sup>+</sup> channel (IRK) | Yes | 0 (0) | 3 (7) | 0 (0) | 1 (2) |

We also detected a magnesium transporter (LtMgT) and an inward rectifier potassium channel (LtIRK) in two *L. tarentolae* BioID datasets, whereas phosphate transporter 91 (LtPho91) and acid phosphatase (LtAP) appeared only in one dataset (**Table 1**). The limited detection of these proteins may reflect spatial separation from the VTC complex within the acidocalcisome membrane; for instance, acid phosphatase is known to localise to the acidocalcisome lumen in *T. brucei* (7), making it unlikely to be directly biotinylated by BirA*.

The complete protein lists from this analysis, including corresponding *L. tarentoale* gene IDs (LtaPh_) and *T. brucei* ortholog accession numbers (Tb927), are provided as searchable supplementary tables (S6-S9 Table).

Further, we assessed the colocalisation of mNG-tagged LtVtc4 with several of these proteins (mCherry (mCh)-tagged LtV-H^+^-ATPase_A, LtPho91, LtCa^2+^-ATPase, LtZnT and LtVtc1) in the acidocalcisome of intact *L. tarentolae* cells using immunofluorescence microscopy; LtVtc1 was the positive control. As expected, LtVtc1 colocalised strongly with LtVtc4, with a Pearson’s correlation coefficient (PCC) of 0.87 (**Fig 3**). LtCa^2+^-ATPase, LtV-H^+^-ATPase_A, LtZnT, and LtPho91 also colocalised with LtVtc4 with PCC of 0.83, 0.87, 0.84, and 0.87, respectively (**Fig 3**), suggesting similar kinds of proximities.

**Figure 3.**
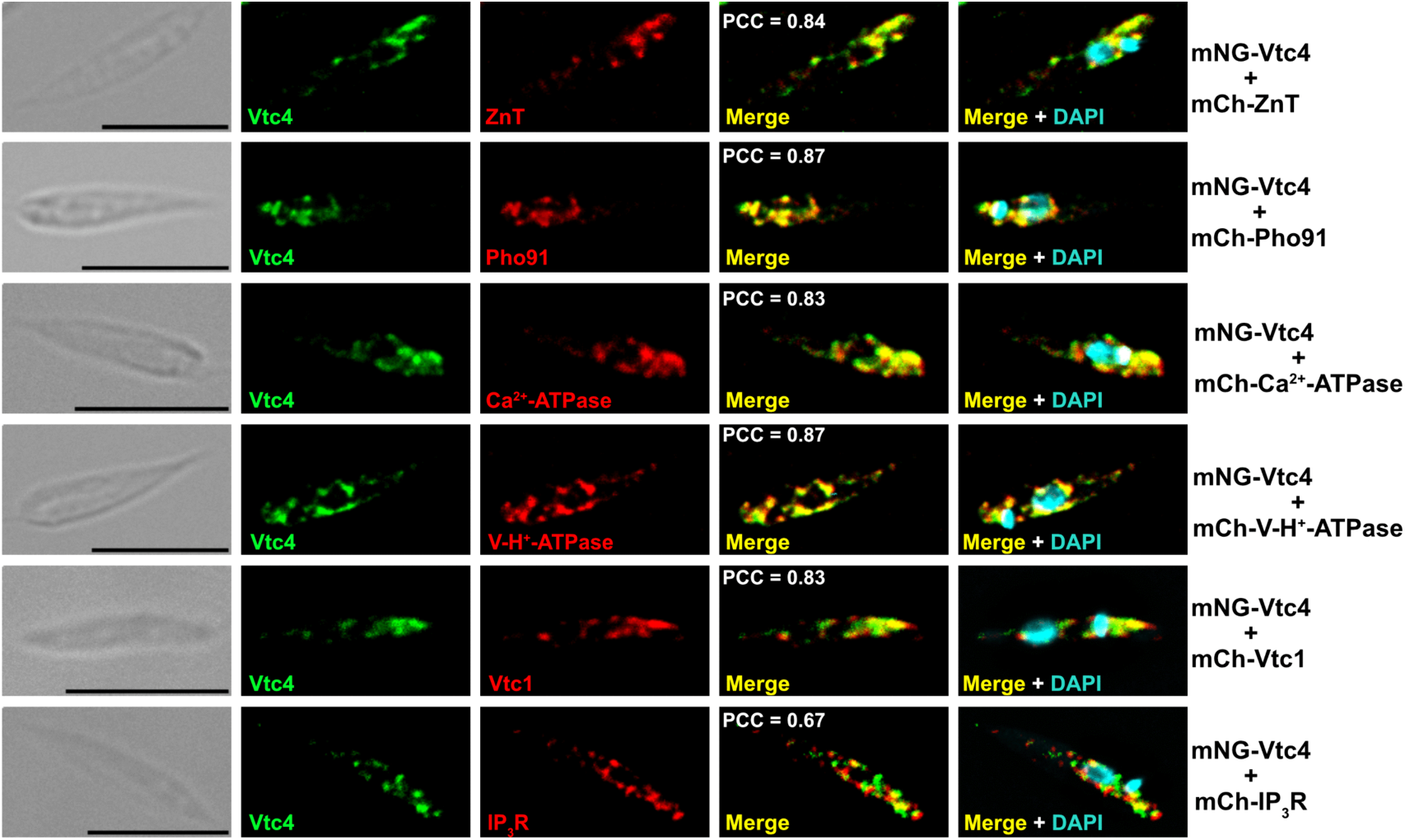
Colocalisation analysis of *L. tarentolae* expressing mNG-LtVtc4 and mCh-tagged by immunofluorescence microscopy. mNG-LtVtc4 colocalises with mCh-LtVtc1 (Pearson’s r = 0.83) as well as with mCh-LtPho91, mCh-LtCa²⁺-ATPase, mCh-LtV-H⁺-ATPase_A, and mCh-LtZnT, showing Pearson’s correlation coefficients of 0.87, 0.83, 0.87, and 0.84, respectively. A lower degree of colocalisation is observed with mCh-IP₃R (r = 0.67). Scale bar: 5 µm.

To test whether these proteins directly interact with the VTC complex, we performed an affinity pulldown assay from cell lysates using ChromoTek mNeonGreen-Trap Agarose beads to capture mNG-tagged LtVtc4. As a positive control, mCh-LtVtc1 was successfully captured with mNG-LtVtc4, as indicated by two strong bands in the bead fraction after washing (**Fig 4A and S5 Fig**). Inositol 1,4,5-triphosphate receptor (IP3R), which was not identified in the BioID datasets, served as a negative control. Although it colocalised with LtVtc4 in the acidocalcisome (**Fig 3**), IP3R was not captured by mNG-Vtc4, confirming the absence of a direct interaction (**Fig 4B and S6 Fig**). Surprisingly, none of the acidocalcisome proteins that colocalised with LtVtc4 were captured in the pulldown assay (**Fig 4B and S6 Fig**), suggesting no direct interaction with LtVtc4. By contrast, LtVtc1 and LtVtc4 appear to form a stable, direct complex interaction (**Fig 4A and S5 Fig**).

**Figure 4.**
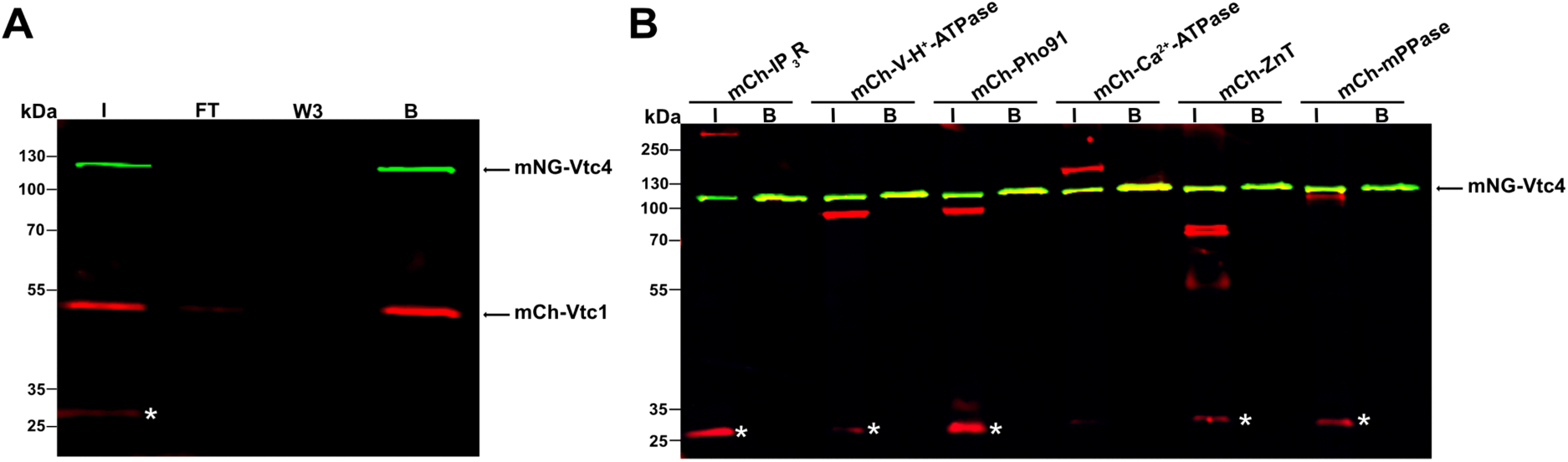
Pulldown assay of acidocalcisome proteins with mNG-LtVtc4. **(A)** mCh-LtVtc1 is captured by mNG-LtVtc4, indicated by two distinct bands in both input and bound fractions. **(B)** No capture is detected for negative control mCh-IP_3_R or for mCh-LtV-H⁺-ATPase_A, mCh-LtPho91, mCh-LtCa²⁺-ATPase, mCh-LtZnT and mCh-LtmPPase, as indicated by the absence of mCh signal in the bound fraction. Lane labels: I – input; FT – flow-through; W3 – third wash; B – bound proteins. Green bands correspond to mNG-LtVtc4; some I and B lines appear yellowish due to partial mNG detection in the 532 nm channel (**see S2 Fig**); red bands correspond to mCh-tagged proteins. The fluorescence signal marked by a white asterisk in the gel represents free mCh tag, visible only in the input due to partial degradation.

In addition to the previously identified acidocalcisome proteins, all datasets contained two novel, putative membrane proteins, LtaPh_2928300 (13.6 kDa) and LtaPh_3617400 (60 kDa), which we named VTC Binding Partner 1 (VBP1) and VTC Binding Partner 2 (VBP2). Further analysis revealed a third hypothetical protein (LtaPh_2928300, 34.4 kDa), present in three out of four datasets (**Table 2**), which we named VTC Binding Partner 3 (VBP3). In *T. brucei*, they are homologous to Tb927.3.3140 (hypothetical protein), Tb927.10.6180 (Fla1-like protein), and Tb927.4.860 (hypothetical protein), respectively.

**Table 2.** Novel VTC binding partners identified by BioID of *L. tarentolae*.

| No. | Gene |  | Protein name | MW (kDa) | Membrane association | Peptide counts (% Coverage) |  |  |  |
| --- | --- | --- | --- | --- | --- | --- | --- | --- | --- |
|  |  |  |  |  |  | DDM |  | FC-12 |  |
|  | *L. tarentolae* | *T. brucei* |  |  |  | BirA-LTVTC1 | BirA-LTVTC4 | BirA-LTVTC1 | BirA-LTVTC4 |
| 1 | LtaPh_2928300 | Tb927.3.3140 | VTC binding partner 1 (VBP1) | 13.6 | Yes | 1 (10) | 2 (25) | 2 (25) | 1 (10) |
| 2 | LtaPh_3617400 | Tb927.10.6180 | VTC binding partner 2 (VBP2) | 60 | Yes | 8 (25) | 5 (14) | 5 (16) | 3 (9) |
| 3 | LtaPh_3438300 | Tb927.4.860 | VTC binding partner 3 (VBP3) | 34.4 | Yes | 3 (13) | 0 (0) | 1 (6) | 1 (6) |

These proteins are unique to kinetoplastids and lack orthologs in other organisms (**S7 Fig**). While the TrypTag database suggests that each ortholog may localise to the acidocalcisome (28), only the LtVPB2 ortholog was found in the *T. brucei* acidocalcisome proteome (7), with experimental studies supporting its acidocalcisomal localisation (49). We used immunofluorescence microscopy to confirm colocalisation of mCh-tagged LtVBP1-3 with mNG-LtVtc4 in acidocalcisomes (**Fig 5**).

**Figure 5.**
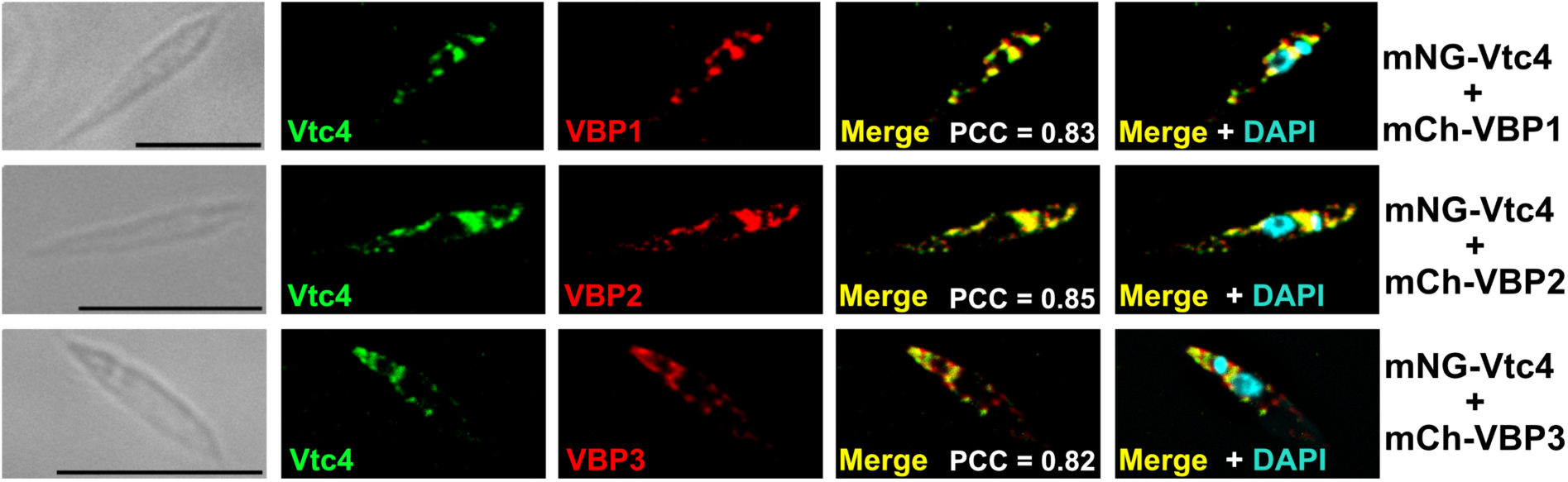
Colocalisation analysis of *L. tarentolae* expressing mNG-LtVtc4 and mCh-tagged LtVBP1, LtVBP2, and LtVBP3 by immunofluorescence microscopy. All three proteins show colocalisation with LtVtc4 in acidocalcisomes. Pearson’s correlation coefficients with LtVtc4 are 0.83 for LtVBP1, 0.85 for LtVBP2, and 0.82 for LtVBP3. Scale bar: 5 µm.

To evaluate direct binding between these proteins and the VTC complex, we conducted pulldown assays as described above, using mNG-LtVtc4 as bait. SDS-PAGE fluorescence visualisation detected mNG-LtVtc4 together with each mCh-LtVBP in the bound fraction, suggesting interaction (**Fig 6A-C and panel A-C in S8 Fig**). However, LtVBP3 bound LtVtc4 more weakly than LtVBP1 or LtVBP2, with a fraction detected in the flowthrough (**Fig 6C and panel C in S8 Fig**). To further validate this interaction, we performed a pulldown assay using mCh-specific beads with mCh-LtVBP3 as bait and confirmed that mNG-LtVtc4 was captured by mCh-LtVBP3 (**Fig 6D and panel D in S8 Fig**), although a significant portion of mNG-LtVtc4 was also detected in the flowthrough.

**Figure 6.**
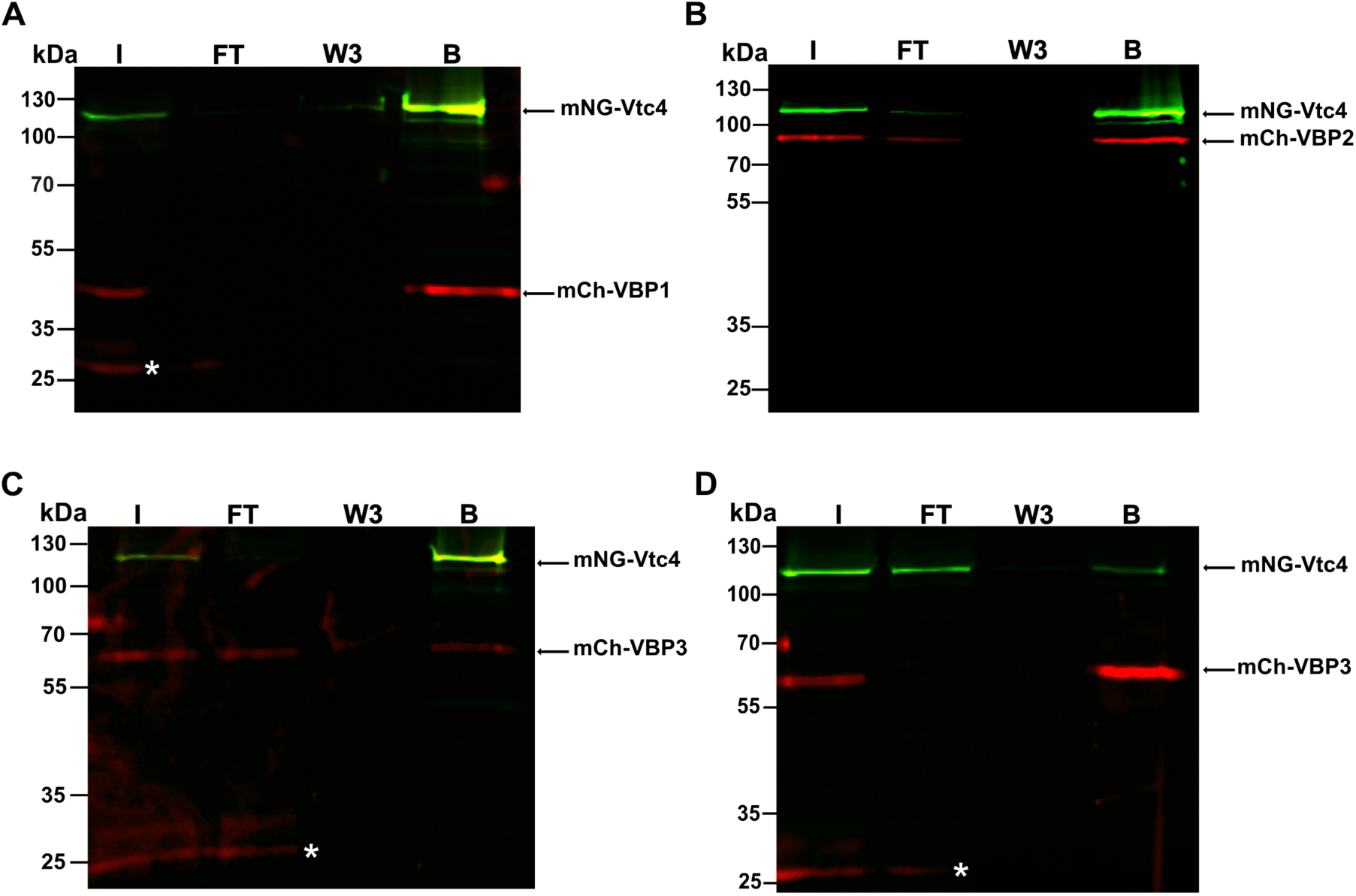
Pulldown assay of novel VTC binding partners with mNG-LtVtc4 in *L. tarentolae.* (A-C) SDS-PAGE with fluorescence detection shows capture of mCh-LtVBP1, mCh-LtVBP2, and mCh-LtVBP3 by mNG-LtVtc4 using mNeonGreen-Trap agarose beads, as indicated by signal in the bound fraction. **(D)** Pulldown of mNG-LtVtc4 with mCh-LtVBP3 using ChromoTek RFP-Trap beads confirms interaction. Lane labels: I – input; FT – flow-through; W3 – third wash; B – bound fraction. Green bands correspond to mNG-LtVtc4; some I and B lines appear as yellowish due to partial detection of the mNG tag in the 532 nm channel (**see S2 Fig**). Red bands correspond to mCherry-tagged proteins. The fluorescence signal marked by a white asterisk in the gel represents free mCh tag, visible only in the input due to partial degradation.

To investigate whether the VTC binding partners influence the localisation of the VTC complex, we analysed LtVtc4 distribution in cells lacking VBP1, VBP2, or VBP3. Knockout transfectant populations without clonal selection (ΔVBP1-Pop, ΔVBP2-Pop, and ΔVBP3-Pop) were generated in a cell line expressing mNG-LtVtc4 (parental cell line) (**S9 Fig**) and examined for acidocalcisome colocalisation with the acidocalcisomal marker mPPase by confocal microscopy (**Fig 7A**). In control parental cells, mNG-LtVtc4 strongly colocalised with mPPase (PCC = 0.74), consistent with its acidocalcisomal localisation (**Fig 7A**). A similar pattern was observed in ΔVBP2-Pop and ΔVBP3-Pop, resulting in PCC of 0.69 and 0.74, respectively (**Fig 7A**). In contrast, ΔVBP1-Pop cells showed reduced colocalisation between LtVtc4 and mPPase (PCC = 0.63). Interestingly, we also observed partial redistribution of LtVtc4 towards the DAPI-stained kinetoplast region (PCC = 0.52) (**Fig 7A**). To quantify these observations, we performed statistical analysis on PCC values obtained from 30 to 50 randomly selected cells using one-way ANOVA. This analysis revealed a strong group effect for acidocalcisomal colocalisation of LtVtc4 (F = 35.23, p = 1.03 × 10^-17^). Tukey’s post hoc test showed that ΔVBP1-Pop differed significantly from the parental cell line, ΔVBP2-Pop and ΔVBP3-Pop (p < 0.0001), whereas no differences were detected among the parental cells, ΔVBP2-Pop, and ΔVBP3-Pop (**Fig 7B**).

**Figure 7.**
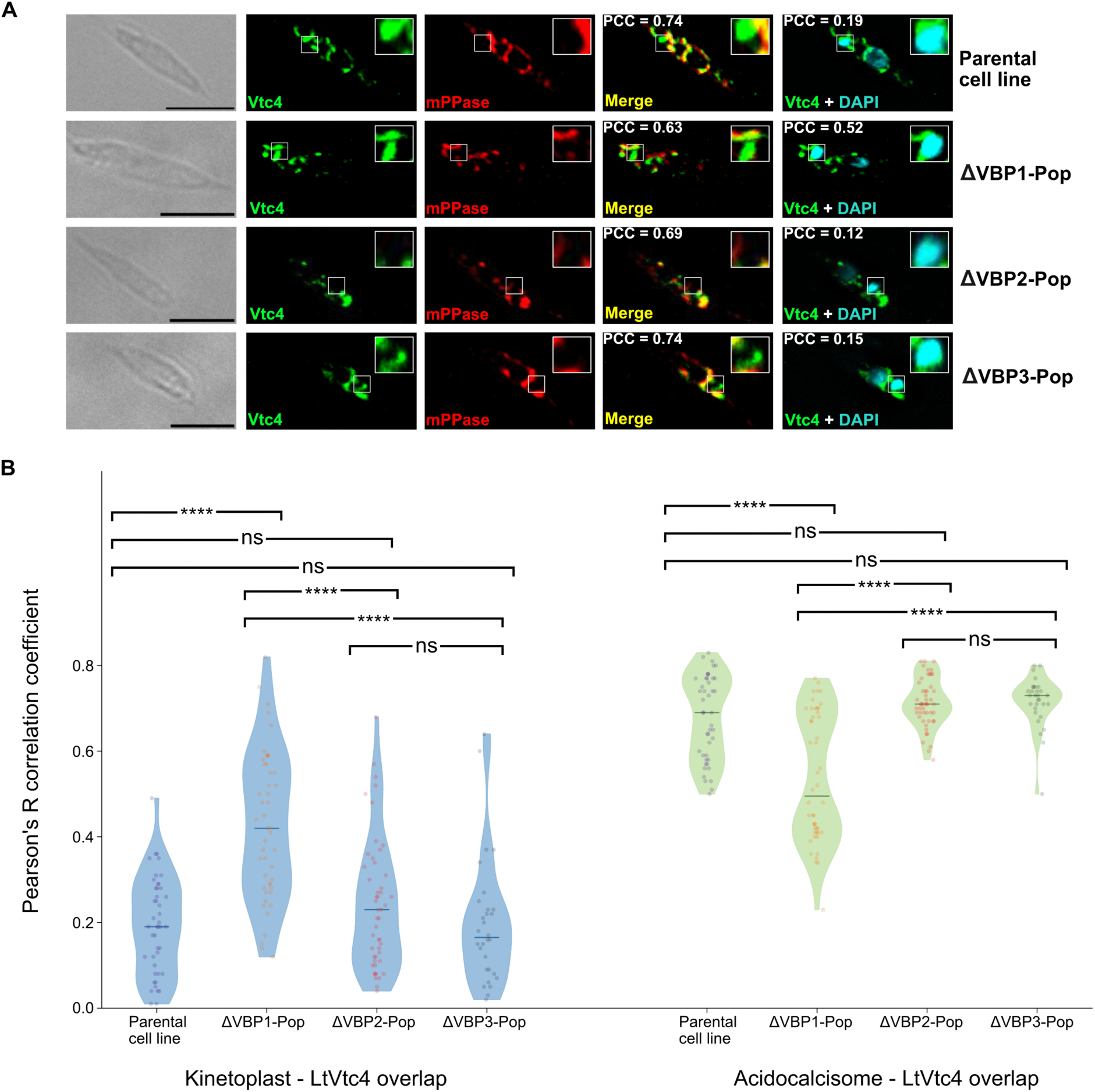
Localisation of mNG-LtVtc4 in VBP knockout population mutants. **(A)** Representative fluorescence microscopy micrographs of *L. tarentolae* cells expressing mNG-LtVtc4 (parental cell line) and in ΔVBP1-Pop, ΔVBP2-Pop and ΔVBP3-Pop. mNG-LtVtc4 colocalises with the acidocalcisome marker mPPase in the parental cell line, ΔVBP2-Pop and ΔVBP3-Pop cells, whereas reduced colocalisation and partial redistribution towards the kinetoplast are observed in ΔVBP1-Pop. Magnified insets highlight the kinetoplast region. Pearson’s correlation coefficients (PCC) for overlapping channels are indicated. Scale bar: 5 µm. **(B)** Quantitative colocalisation analysis of LtVtc4 localisation. Violin plots show PCC values calculated from 30 to 50 randomly selected cells for each strain. Blue violin plots represent LtVtc4 colocalisation with kinetoplast DNA stained with DAPI; green violin plots represent LtVtc4 colocalisation with the acidocalcisome marker mPPase. Each dot represents one analysed cell. Statistical significance was assessed using one-way ANOVA followed by Tukey’s post hoc test. ****p < 0.0001.

Consistent with the immunofluorescence microscopy observations, analysis of LtVtc4 colocalisation with DAPI-stained kinetoplast DNA also showed a group effect (F = 25.04, p = 1.92 × 10^-13^), with ΔVBP1-Pop showing higher correlation values than the parental cell line and the other knockout transfectant populations (**Fig 7B**). These results indicate that VBP1 affects the correct localisation of LtVtc4 to the acidocalcisome, while deletion of VBP2 or VBP3 does not.

### Predicted structures of the VTC complex components and their binding partners support complex assembly

Following confirmation of direct interaction by pulldown assays, we used AlphaFold3 (41) to predict the structures of LtVtc1, LtVtc4, and their binding partners, and assessed the potential for complex formation. The transmembrane topology and orientation of each protein were predicted using MEMSAT-SVM *via* the PSIPRED server (44, 45). AlphaFold3 predicted well-folded structures for all proteins, with most residues showing high-confidence pLDDT scores (pLDDT > 70), indicating good confidence in the model (**Fig 8A**). The C-terminal TMHs of both LtVtc1 and LtVtc4 have an average pLDDT above 70. In contrast, the N-terminal region of LtVtc1 (aa1-59) is unstructured, with pLDDT scores below 50, indicating very low confidence in the model. The N-terminal cytoplasmic region of LtVtc4, containing an SPX domain and a CD domain, is well structured with high pLDDT scores (**Fig 8A**), except in the loops that connect the domains. Unlike LtVtc1 and LtVtc4, all VBPs have only a single-pass TMH, at the N-terminus for LtVBP1 and LtVBP2 and at the C-terminus for LtVBP3 (**Fig 8A**). On the lumenal side, LtVBP1 has two β-hairpin motifs; LtVBP2 contains a β-propeller domain composed of six four-stranded β-sheets; and LtVBP3 consists of a C-terminal domain structurally similar to LtVtc4-CD with an RMSD of 5.85 Å (**Fig 8B**) and to the inorganic polyphosphatase from *Escherichia coli* (EcygiF) (53) with an RMSD of 4.13 Å (**Fig 8C**), despite very low sequence identity – 10.6% with LtVtc4-CD and 15.4% with EcygiF-NTD, respectively (**S10 Fig**).

**Figure 8.**
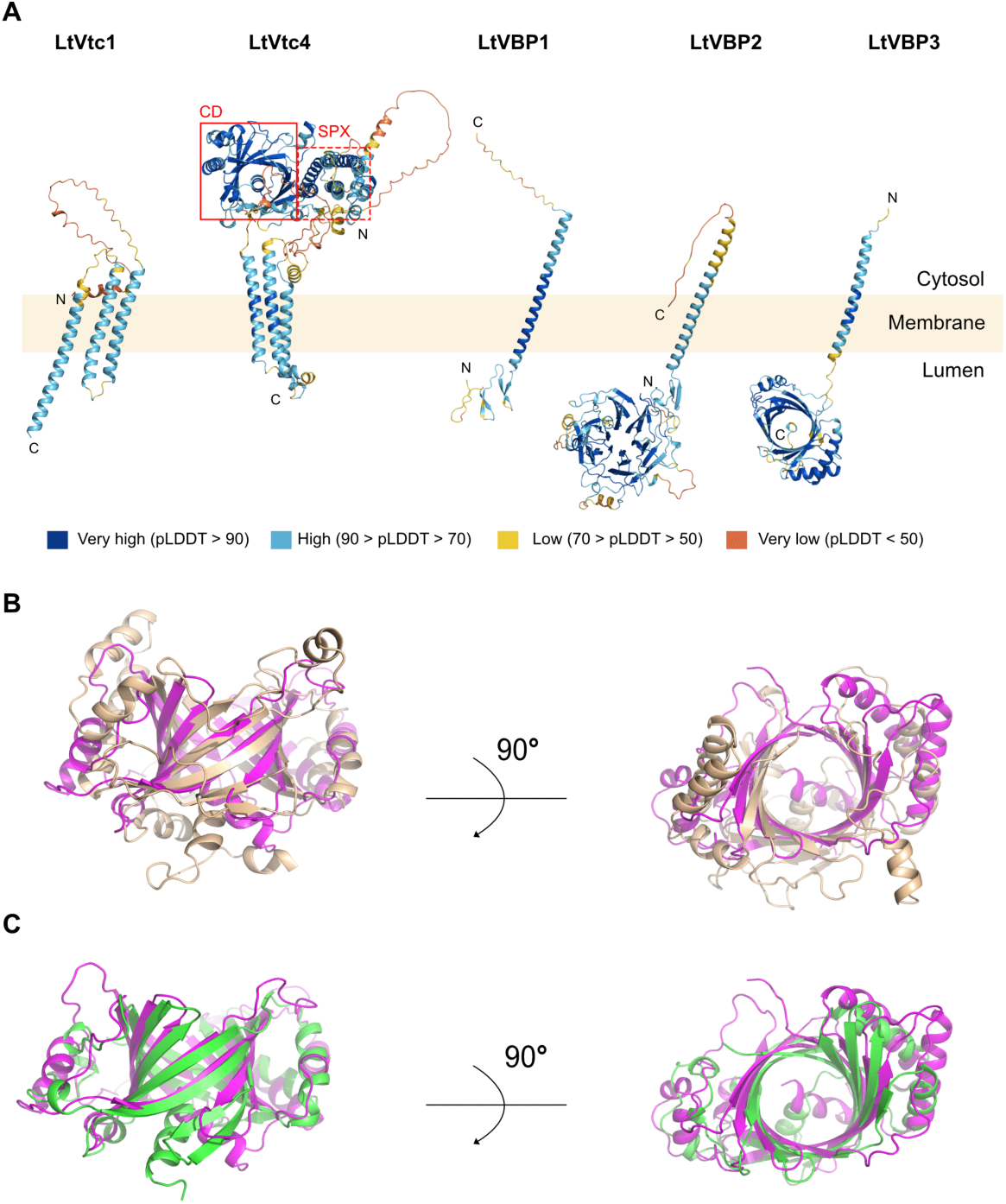
Novel VTC complex interactors. **(A)** AlphaFold3 model of LtVtc1, LtVtc4, LtVBP1, LtVBP2, and LtVBP3. Transmembrane topology and orientation were predicted using MEMSAT-SVM (45). The colour is based on the pLDDT colouring scheme. N: N-terminal residue, C: C-terminal residue. Red square: CD domain of Vtc4 and dashed-red square: SPX domain of Vtc4. **(B)** The predicted structure of the C-terminal domain (CTD) of LtVBP3 (magenta) superimposes on the catalytic domain of LtVtc4 (wheat) with an RMSD of 5.85 Å. **(C)** The predicted structure of the C-terminus domain of LtVBP3 (magenta) on the structure of inorganic polyphosphatase from Escherichia coli (EcygiF) (PDB: 5a60) (green) with an RMSD of 4.13 Å.

To analyse complex formation between the VTC components with LtVBPs, we used AlphaFold3 to predict all pairwise interactions among the five proteins and evaluated the inter-chain predicted TM-score (ipTM) (54) for each combination (**panel A in S11 Fig**). The LtVtc1-LtVtc4 complex was used as a positive control, as this interaction has been structurally confirmed by the yeast VTC complex (15). Although the ipTM score for the LtVtc1-LtVtc4 is only 0.62, which is relatively low, the predicted aligned error (PAE) plot reveals consistent low-error regions between subunits, indicating a good intermolecular structural correlation (**panel B in S11 Fig**). Among the predicted complexes, LtVtc4-LtVBP1, LtVtc4-LtVBP2, and LtVBP1-LtVBP3 have ipTM scores of 0.50, 0.52, and 0.66, respectively (**panel A in S11 Fig**). To further assess these interactions, we calculated the actifpTM (actual interface pTM) scores, which focus specifically at predicted interfacial residues (43). The results show that the actifpTM scores of LtVtc4 in complexes with LtVtc1, LtVBP1, LtVBP2, and LtVBP1 with LtVBP3 are 0.93, 0.91, 0.92, and 0.96, respectively (**panel A in S11 Fig**), which indicates strong confidence in complex formation. Interestingly, the interacting sites of the complexes align with the transmembrane orientation of each protein predicted by MEMSAT-SVM (**Fig 8A**).

Based on the predicted structures, and consistent with the yeast VTC complex structure (15), LtVtc4 forms a complex with two LtVtc1 subunits *via* their transmembrane helices (TMHs) at two distinct interfaces: interface 1 pairs TMH1/TMH3 of the first LtVtc1 with TMH1/TMH2 of LtVtc4, whereas interface 2 pairs TMH1/TMH2 of the second LtVtc1 to TMH1/TMH3 of LtVtc4 (**Fig 9A and panel C in S11 Fig**). Although the yeast complex also forms Vtc1-Vtc1 interactions, the predicted LtVtc1–LtVtc1 homodimer scores were low (ipTM = 0.41; actifpTM = 0.40), suggesting a possible false-negative prediction (**panel A in S11 Fig**). Despite that limitation, the model indicates two additional contact sites on LtVtc4. A cytoplasmic helix-turn-helix of LtVtc4-CD domain engages a short helix in LtVBP1, while the lumenal end of TMH3 contacts the junction between the β-propeller and single TMH of LtVBP2, respectively (**Fig 9B-C and panel D-E in S11 Fig**). Both interfaces are dominated by hydrophobic contacts with a few hydrogen bonds and salt bridges. No direct interface is predicted for LtVBP3; its low ipTM/actifpTM values agree with a model in which LtVBP3 binds LtVBP1 *via* its single TMH and adjacent loop, rather than LtVtc4 itself, forming a β-sheet with the first β-hairpin of LtVBP1 (**Fig 9D and panel F in S11 Fig**). This suggests that, unlike LtVBP1 and LtVBP2, which bind directly to Vtc4, LtVBP3 binds indirectly to the VTC complex *via* its interaction with LtVBP1.

**Figure 9.**
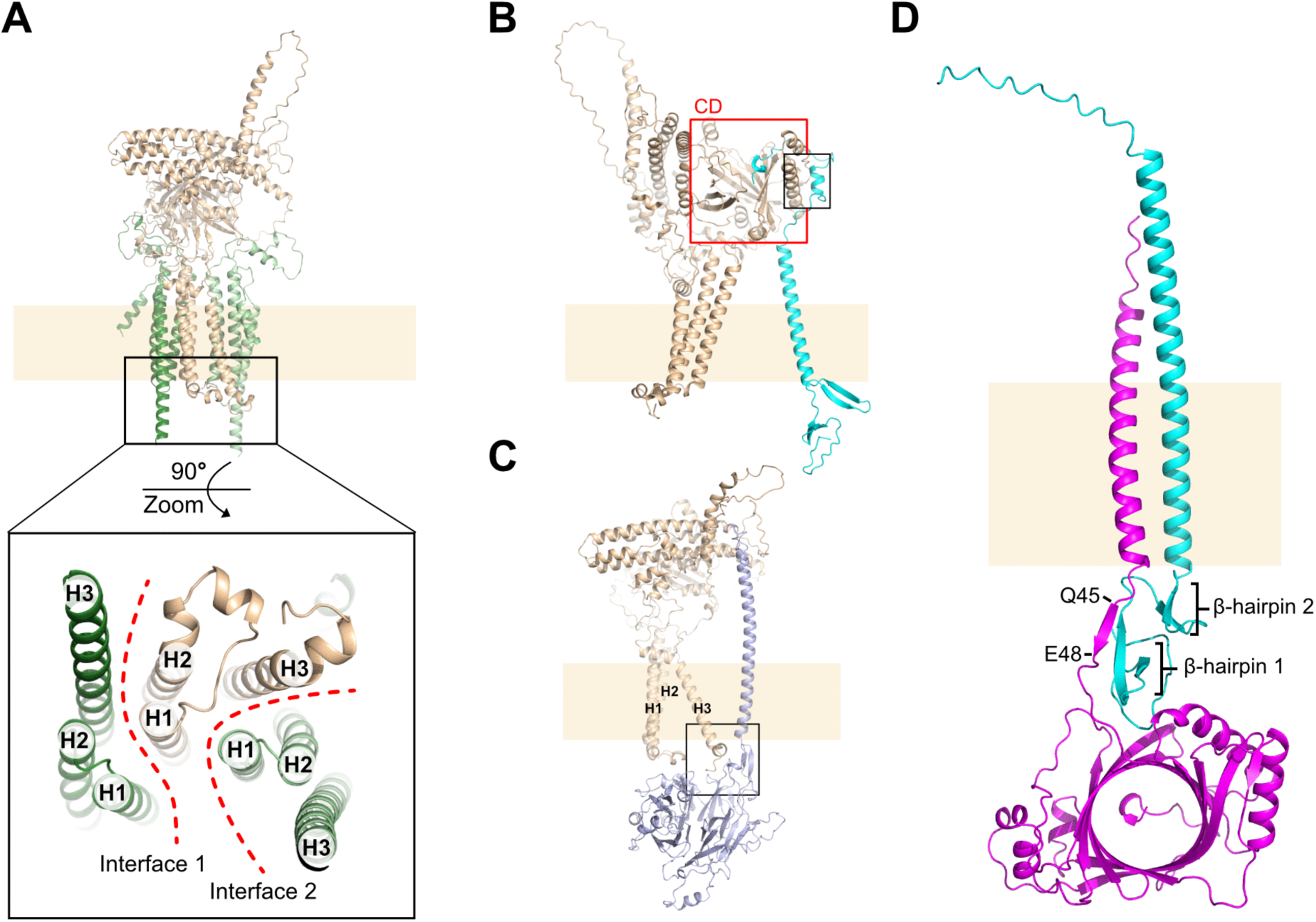
AlphaFold3 prediction of the complex formed by VTC components and VBPs. **(A)** Complex prediction of two LtVtc1 (green) with one LtVtc4 (wheat). The zoomed-in inset shows the arrangement of TMHs of each LtVtc1 and LtVtc4 viewed from the lumenal side. H1, 2, 3 = TMH1, 2, and 3, respectively. Red-dashed lines are interfaces separating TMH of LtVtc1 and LtVtc4. **(B)** Complex prediction of LtVtc4 (wheat) with LtVBP1 (cyan). Red rectangle: CD domain of LtVtc4. Black rectangle: Interaction region between LtVtc4 and LtVBP1. **(C)** Complex prediction of LtVtc4 (wheat) with LtVBP2 (light blue). Black rectangle: Interaction region between the TMH3 of LtVtc4 with LtVBP2. **(D)** Complex prediction of LtVBP1 (cyan) with LtVBP3 (magenta).

## Discussion

To date, over 30 proteins have been identified in kinetoplastid acidocalcisomes, primarily through proteomic studies in *T. brucei* (6), with more limited data from *T. cruzi* (55) and *L. donovani* (10), as well as localisation data from the TrypTag.org project (28, 49, 56). Among these proteins, the VTC complex is one of the key components of kinetoplastid acidocalcisome function, but its composition remains elusive. In yeast, the complex consists of five subunits (15), but only Vtc1 and Vtc4 are conserved and are found in kinetoplastids. This highlights a major gap in our understanding of how the VTC complex has been remodelled in kinetoplastids and whether its function is supported by additional lineage-specific protein components. To address this gap, we used the BioID method in *L. tarentolae* to identify novel VTC-interacting proteins in the acidocalcisomes.

In yeast, the VTC complex comprises three Vtc1, one Vtc4, and one Vtc2/3 subunit (15). Here, we initially sought VTC subunits in *L. tarentolae* with structural similarity to Vtc2 or Vtc3, despite low or no sequence identity. However, we did not observe any Vtc2- or Vtc3-like subunits in our BioID data. Instead, we identified three unique single-pass TMH proteins (LtVBP1-3) conserved only in kinetoplastids (**S7 Fig**) and confirmed their colocalisation with LtVtc4 to acidocalcisomes (**Fig 5**). Interestingly, population knockout of LtVBP1, but not LtVBP2 or LtVBP3, altered the localisation of LtVtc4, leading to its partial redistribution to the kinetoplast (**Fig 7**). This observation suggests that LtVBP1 may contribute to the proper targeting or stabilisation of the VTC complex within acidocalcisomes. Pulldown assays (**Fig 6A–D**) and AlphaFold3 predictions (**Fig 9**) further validated direct interactions among these proteins (**Fig 10**).

**Figure 10.**
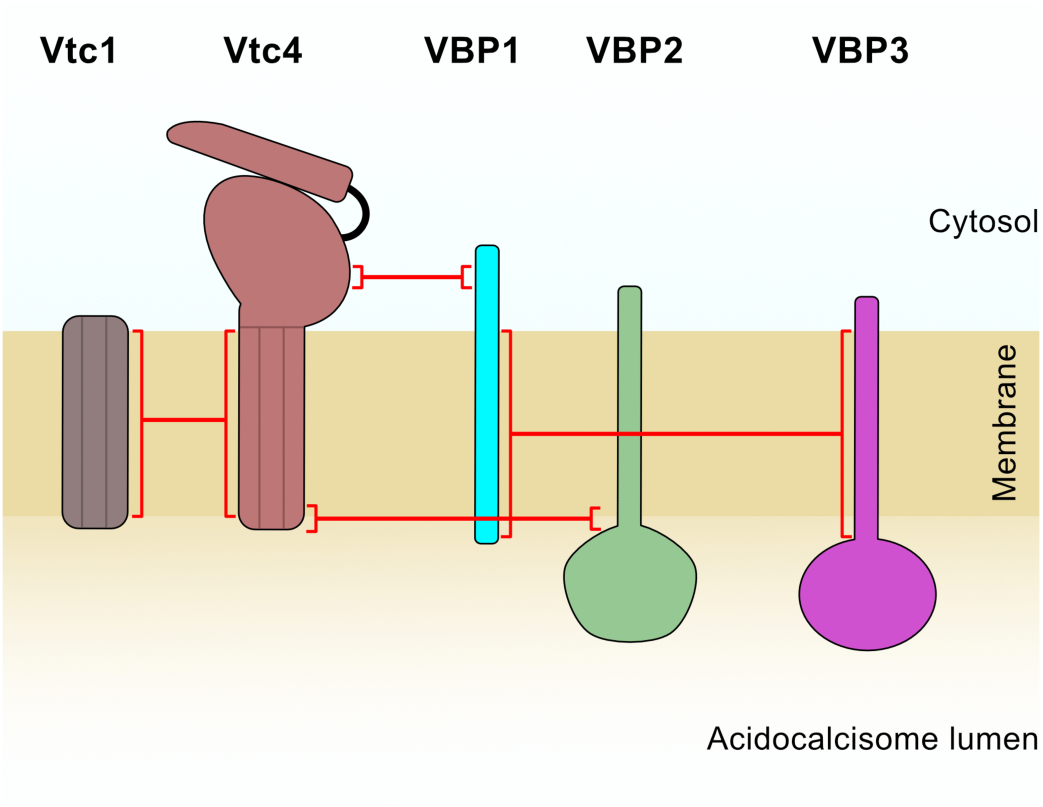
Schematic interaction map of the VTC complex with the three novel VTC-binding proteins (VBP1-3). Red lines mark the regions where physical interactions between VTC subunits and VBPs were predicted based on AlphaFold3. The cytosolic face is oriented upward and the acidocalcisome lumen downward.

Based on the AlphaFold3 prediction, the C-terminal region of LtVBP1 interacts with LtVtc4-CD (**Fig 9B**) in the cytoplasmic space, while its single-pass TMH and N-terminal β-hairpin motifs interact with the single-pass TMH of LtVBP3 (**Fig 9D**). No high-confidence interactions were predicted between LtVBP3 and either LtVtc1 or LtVtc4 (**panel A in S11 Fig**), although this could represent a false negative, as seen with the Vtc1-Vtc1 complex (**panel A in S11 Fig**). Interestingly, the LtVBP3-CTD is structurally similar to LtVtc4-CD (**Fig 8B**) and EcygiF (**Fig 8C**) despite having very low sequence identity and a currently unknown function.

LtVBP2, on the other hand, is predicted to interact with the C-terminal region of LtVtc4 *via* the neck that links its single-pass transmembrane helix (TMH) to the β-propeller domain (**Fig 9C**). The presence of such a domain in the putative VTC complex is a surprise. However, β-propeller domains support diverse cellular functions: four-bladed propellers frequently mediate transport; five-bladed variants can act as transferases or hydrolases; and larger, six- to eight-bladed propellers are often associated with structural and signaling roles (57). Among parasites, such β–propeller–containing proteins participate in a range of processes. For example, in *T. brucei*, oligopeptidase B uses a β-propeller for substrate gating and virulence (58); in *Plasmodium falciparum*, the β-propeller domain of K13 is linked to artemisinin resistance (59); and in trypanosomatids, WD-repeat β-propeller proteins contribute to cell cycle control and signaling (60). Given that LtVBP2 directly interacts with the VTC complex and contains a six-bladed β-propeller, it may be involved in structural or signaling functions within the complex, consistent with roles attributed to larger β-propellers (57).

During revision of this manuscript, a recent study characterising the VTC complex in *T. cruzi* reported TcVtc6 (61), a homolog of LtVBP3 identified in this study, as an important factor for polyP synthesis, differentiation, and parasite infectivity. The proteomic dataset also included homologs of LtVBP1 and LtVBP2, although these proteins were not further characterised as potential VTC-associated factors (61). These findings independently support the presence of additional conserved kinetoplastid-specific proteins associated with the VTC complex. While deletion of LtVBP3 did not appear to affect VTC complex localisation in *L. tarentolae*, the infectivity phenotype reported in *T. cruzi* (61) suggests that this protein may also contribute to parasite fitness or host interaction processes. It is therefore possible that VBP3 also plays a role in infectivity-related functions in pathogenic *Leishmania* species.

In summary, our study provides a coherent interaction framework that expands the known protein environment of the VTC complex in acidocalcisomes. The BioID data establish a molecular context in which the kinetoplastid VTC complex operates close to several acidocalcisome membrane proteins, including LtPPase, LtCa²⁺-ATPase, LtZnT, LtPAT2, and subunits of the LtV-H⁺-ATPase. These proteins were consistently detected across BioID datasets (**Table 1**), suggesting a reproducible pattern of spatial association with the VTC complex within the acidocalcisome membrane, despite the absence of direct interactions as assessed by pulldown assays (**Fig 4B**). Such proximity may reflect indirect or coordinated roles in polyphosphate synthesis, regulation, or storage within the organelle.

Moreover, we identified VBP1–3 as the first VTC-associated proteins beyond Vtc1 and Vtc4 in kinetoplastids, exemplified by *L. tarentolae*, with recent findings in *T. cruzi* further supporting the conserved association of these proteins with the kinetoplastid VTC complex (61). Confocal microscopy and colocalisation analyses suggest that LtVBP1 may contribute to the stabilisation of the complex and directing acidocalcisome localisation, whereas LtVBP2, with its predicted six-bladed β-propeller fold, could provide structural support or mediate signalling functions that influence phosphate homeostasis. LtVBP3 appears to interact more weakly, which may indicate a transient or regulated association. These findings clearly suggest a kinetoplastid-specific adaptation of the VTC complex, distinct from the architecture described in yeast. By defining the protein interaction partners of the VTC complex, this work establishes a foundation for future experimental studies aimed at resolving the composition and organisation of the kinetoplastid VTC complex. However, the stoichiometry and overall arrangement remain unresolved. High-resolution structural approaches, such as cryo-electron microscopy, combined with targeted genetic and biochemical experiments, will be essential to clarify the roles of VBPs and determine whether they function as integral components of the VTC machinery or as context-dependent interactors.

## Acknowledgements

We thank Eva Gluenz and Tom Beneke for providing plasmids for CRISPR/Cas9 editing in *L. tarentolae*. We acknowledge Rabah Soliymani and the Meilahti Clinical Proteomics Core facility, supported by Biocenter Finland, for technical assistance and mass spectrometry analyses. Imaging was performed at the Light Microscopy Unit, Institute of Biotechnology, supported by HiLIFE and Biocenter Finland.

