## Supporting information for "Novel Binding Partners of the Vacuolar Transporter Chaperone (VTC) complex in Acidocalcisomes of *Leishmania tarentolae*"

**S1 Fig. Acidocalcisome isolation of the BirA* tagged LtVtc1 and Vtc4 from *L. tarentolae*.**

**S2 Fig. SDS-PAGE analysis demonstrating fluorescence signal overlap in gels visualised using the Sapphire Biomolecular Imager.**

**S3 Fig. Purification of LtVtc4-mNG using anti-c-Myc affinity resin and polyP activity assay by urea-PAGE.**

**S4 Fig. Gene ontology of the total protein hits (1240 unique proteins) from the combined VTC BioID results.**

**S5 Fig. Pulldown assays evaluating direct interactions between mNG-LtVtc4 and mCh-LtVtc1.**

**S6 Fig. Pulldown assays evaluating direct interactions between mNG-LtVtc4 and acidocalcisomal membrane proteins tagged with mCh.**

**S7 Fig. BLASTP search in UniprotKB and SwissProt Reference proteins database for (A) LtVBP1, (B) LtVPB2, and (C) LtVBP3, showing protein conservation only in kinetoplastids.**

**S8 Fig. Pulldown assays evaluating direct interactions between mNG-LtVtc4 and novel VTC binding partners (VBPs1-3) tagged with mCh.**

**S9 Fig. Strategy used for VBP knockout in *Leishmania tarentolae*.**

**S10 Fig. Sequence alignment between LtVtc4-CD, LtVBP3-CTD, and EcygiF-NTD shows very low sequence identity: 10.6% between LtVtc4-CD and LtVBP3-CTD and 15.4% between LtVBP3-CTD and EcygiF-NTD.**

**S11 Fig. AlphaFold3 complex structure prediction of the VTC complex with its binding proteins.**

**S1 Table. List of primers used for PCR amplification and confirmation of tagging and knockout cassettes, and corresponding sgRNAs.**

**S2 Table. Proteins identified by BioID affinity purification followed by LC–MS analysis of BirA*–LtVtc1 samples solubilised with 2% DDM.**

**S3 Table. Proteins identified by BioID affinity purification followed by LC–MS analysis of BirA*–LtVtc1 samples solubilised with 2% FC-12.**

**S4 Table. Proteins identified by BioID affinity purification followed by LC–MS analysis of BirA*–LtVtc4 samples solubilised with 2% DDM.**

**S5 Table. Proteins identified by BioID affinity purification followed by LC–MS analysis of BirA*–LtVtc4 samples solubilised with 2% FC-12.**

**S6 Table. Proteins with predicted glycosome localisation.**

**S7 Table. Proteins predicted to localise to RNA granules/polysomes.**

**S8 Table. Proteins predicted to be polyphosphorylated.**

**S9 Table. Total protein possibly located in the small cytoplasmic organelles based on the TrypTag server.**

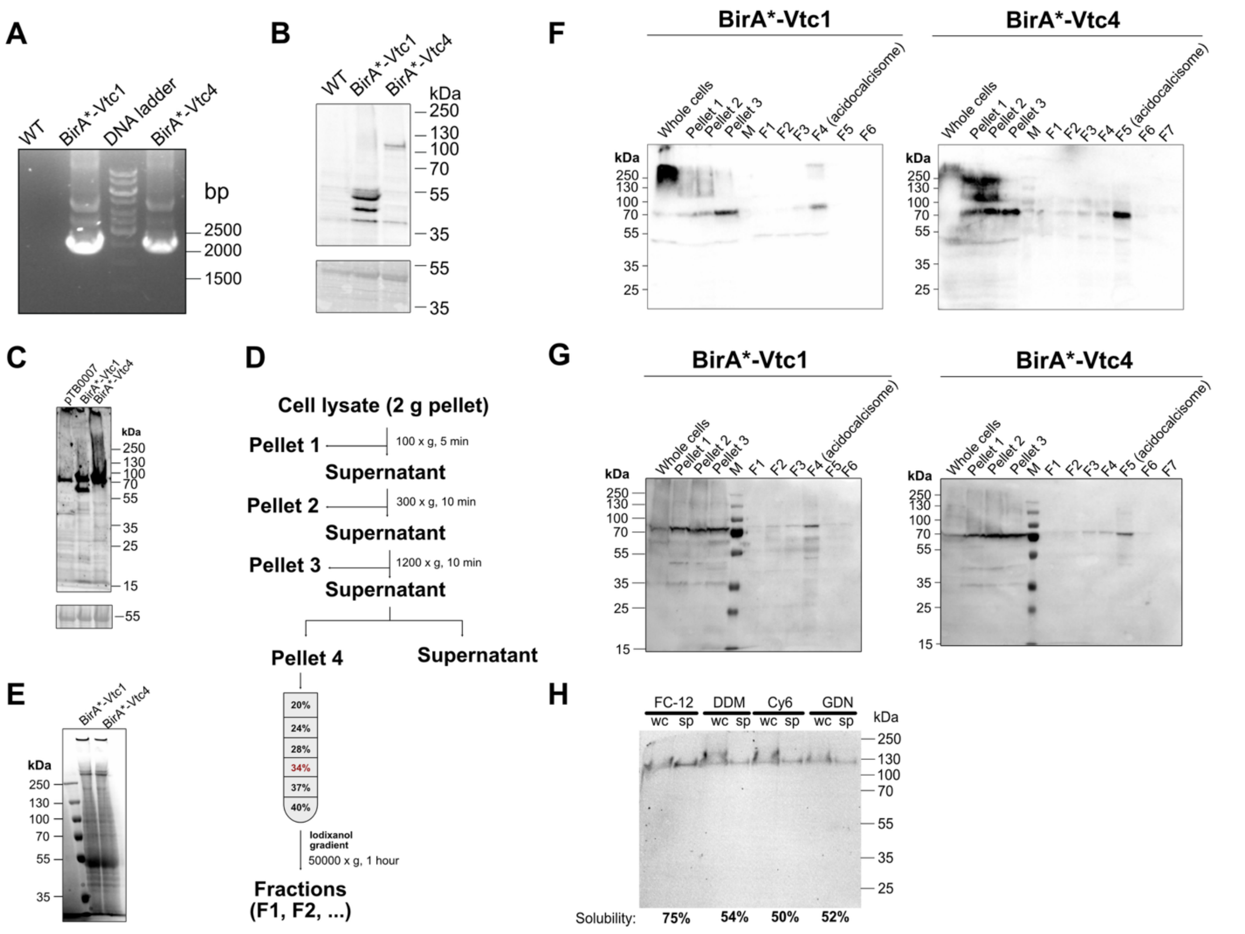


**S1 Fig. Acidocalcisome isolation of the BirA* tagged LtVtc1 and Vtc4 from *L. tarentolae*.**

**(A)** PCR product of BirA*-tagging cassette integration in LtVtc1 and LtVtc4 shows clear bands around 2200 bp for both, while no bands are seen in the wt lane. **(B)** Western blot detection of BirA*-tagged LtVtc1/LtVtc4 expressed in *L. tarentolae* using anti-Myc. Clear bands are observed in LtVtc1 and LtVtc4 lanes and absent in WT. Ponceau S staining of the membrane is shown below as loading control. **(C)** Western blot of cells expressing BirA* tagged LtVtc1 and Vtc4 after being treated with 50 μM biotin for 40h. pTB007: cells containing the pTB007 plasmid as a negative control. Ponceau S staining of the membrane is shown below as a loading control. **(D)** Acidocalcisome fractionation protocol. **(E)**

Acidocalcisome fraction after iodixanol gradient ultracentrifugation. **(F)** Western blot of the iodixanol gradient fractionation using an anti-TmPPase antibody. **(G)** Western blot of the iodixanol gradient fractionation using Avidin-HRP. **(H)** SDS-PAGE showing the mNG fluorescence signal from the detergent solubilisation assay of mNG-LtVtc4 using FC-12 (Fos-choline-12), DDM (*n*-dodecyl-β-maltoside), Cy6 (Cymal-6), and GDN (Glyco-Diosgenin). wc= whole cell sample, sp= supernatant after centrifugation.

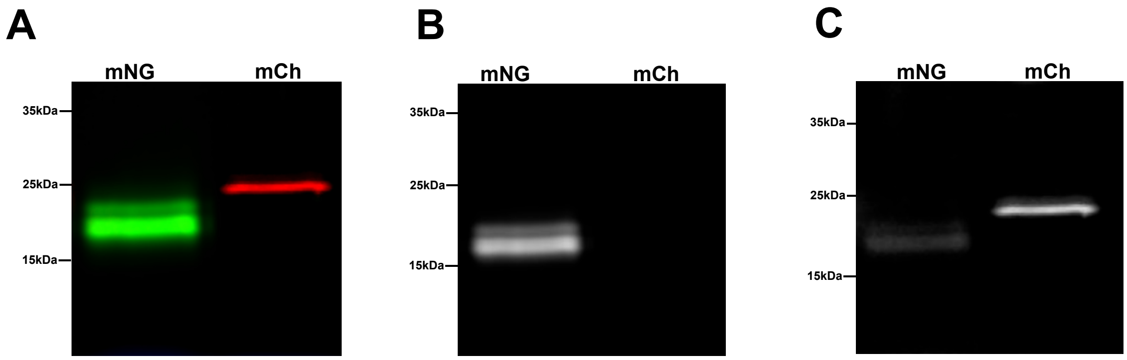


**Supplementary Figure 2.** SDS-PAGE analysis demonstrating green (488 nm) and red (532 nm) fluorescence signals in gels visualized using the Sapphire Biomolecular Imager. **(A)** Merged visualisation of purified mNG and mCh tags. **(B)** Signal of the purified mNG tag alone, displaying exclusively green fluorescence in the 488 nm channel. **(C)** Signal of the purified mCh tag alone in the 532 nm channel, showing strong red fluorescence and weaker green fluorescence due to spectral bleed-through from weak excitation at 532 nm and the emission tail into the red detection range. As a result, merged images may display yellowish bands, depending on the relative green signal intensity.

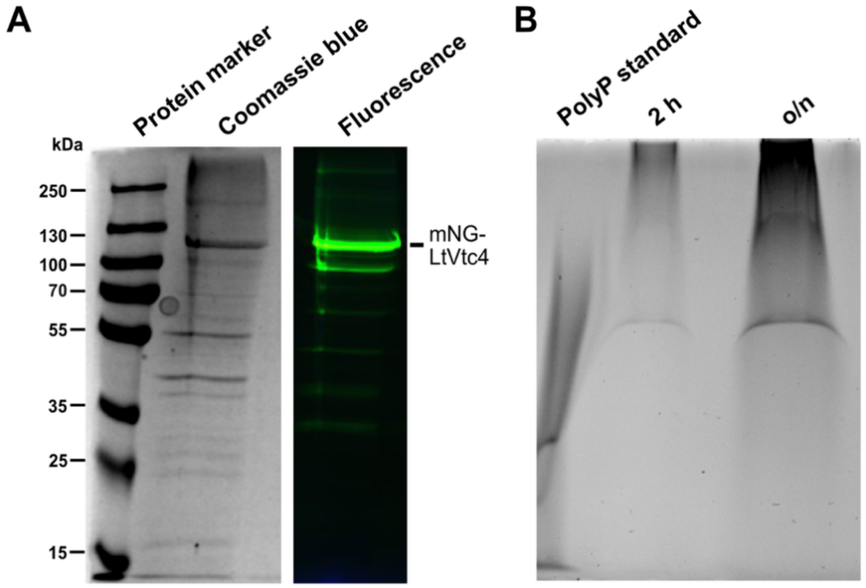


**S3 Fig. Purification of LtVtc4-mNG using anti-c-Myc affinity resin and polyP activity assay by urea-PAGE. (A)** SDS–PAGE analysis of the purified VTC complex visualised by Coomassie Brilliant Blue staining and in-gel fluorescence showing mNG-LtVtc4 signal. **(B)** Urea–PAGE analysis of polyP synthesis activity of the purified VTC complex. The image shows 0.1 µg of polyP standard and polyP produced by the purified VTC complex after incubation with ATP and $IP_6$ at 30 °C for 2 h and overnight.

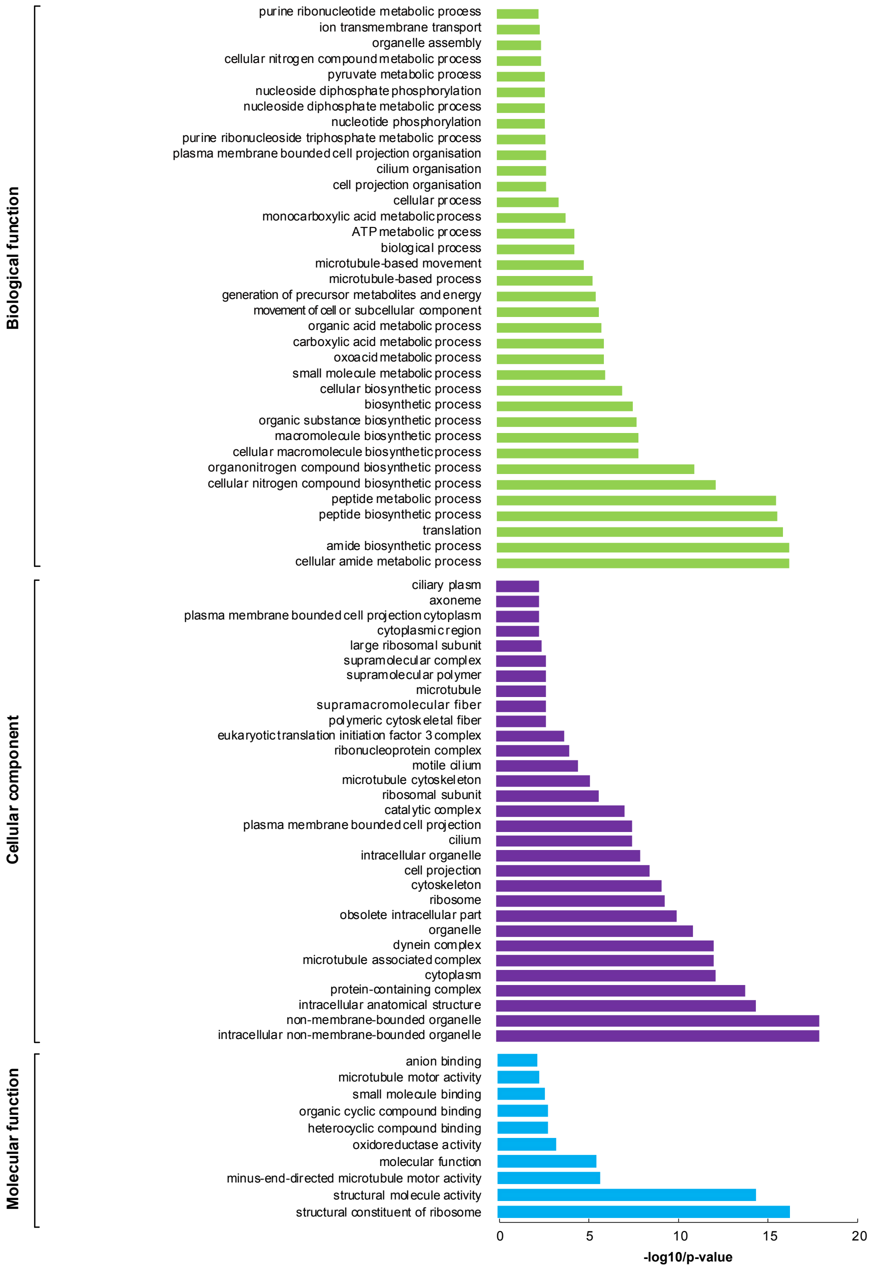


**Supplementary Figure 4**. Gene ontology of the total protein hits (1240 unique proteins) from the combined VTC BioID results.

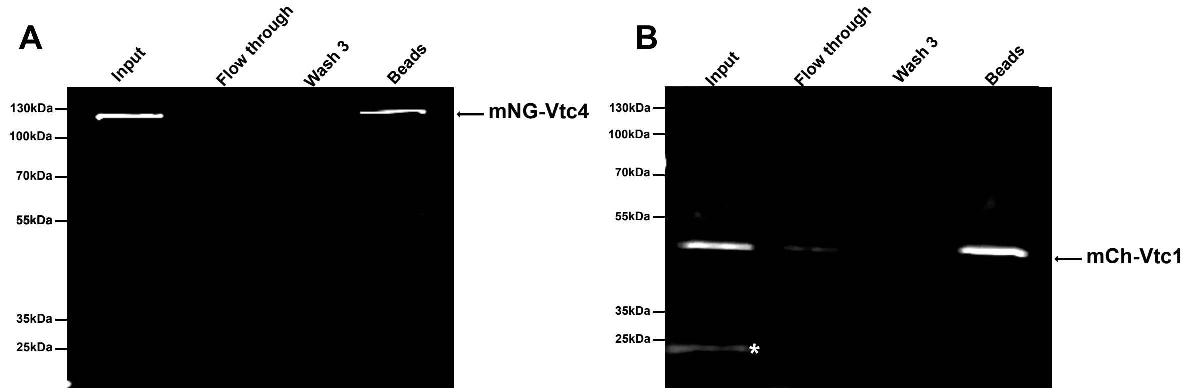


**Supplementary Figure 5.** Pulldown assays evaluating direct interactions between mNG-LtVtc4 and mCh-LtVtc1. **(A)** Detection of mNG-LtVtc4 signal in the green fluorescence channel (488 nm), indicating the presence of LtVtc4 in the assay. **(B)** Detection of mCh-LtVtc1 signal in the red fluorescence channel (532 nm). The mNG signal may also be partially visible in this channel, depending on its relative intensity, which result in yellow bands reflecting overlapping fluorescence in the merged images.

The fluorescence signal marked by a white asterisk in the gel corresponds to the mCh tag alone, visible only in the input due to partial degradation.

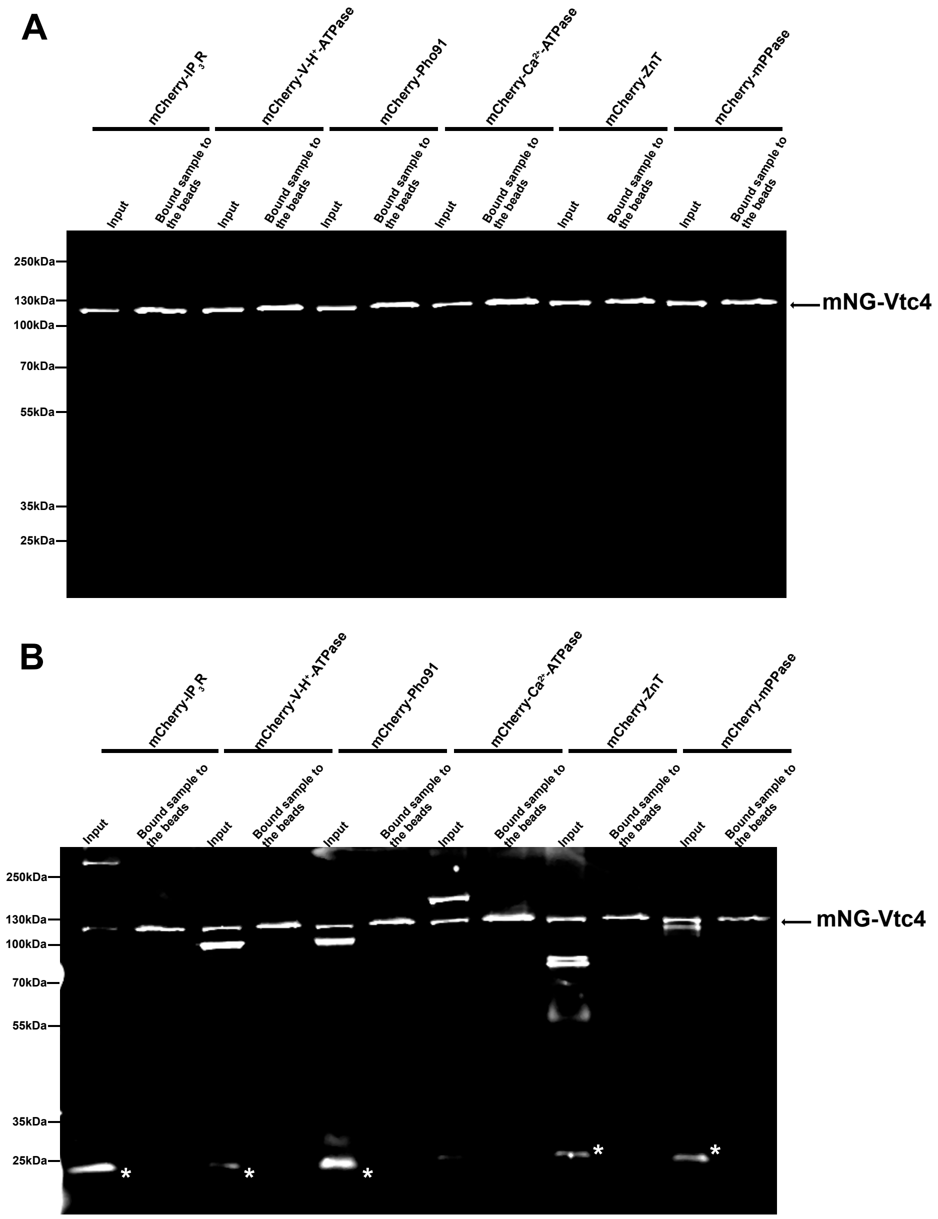


**Supplementary Figure 6.** Pulldown assays evaluating direct interactions between mNG-LtVtc4 and acidocalcisomal membrane proteins tagged with mCh. **(A)** Detection of mNG-LtVtc4 signal in the green fluorescence channel (488 nm), indicating the presence of Vtc4 in the assay. **(B)** Detection of mCh-tagged acidocalcisomal proteins in the red fluorescence channel (532 nm). The mNG signal may also be partially visible in this channel, depending on its relative intensity, which result in yellow bands reflecting overlapping fluorescence in the merged images. The fluorescence signal marked by a white asterisk in the gel corresponds to the mCh tag alone, visible only in the input due to partial degradation.

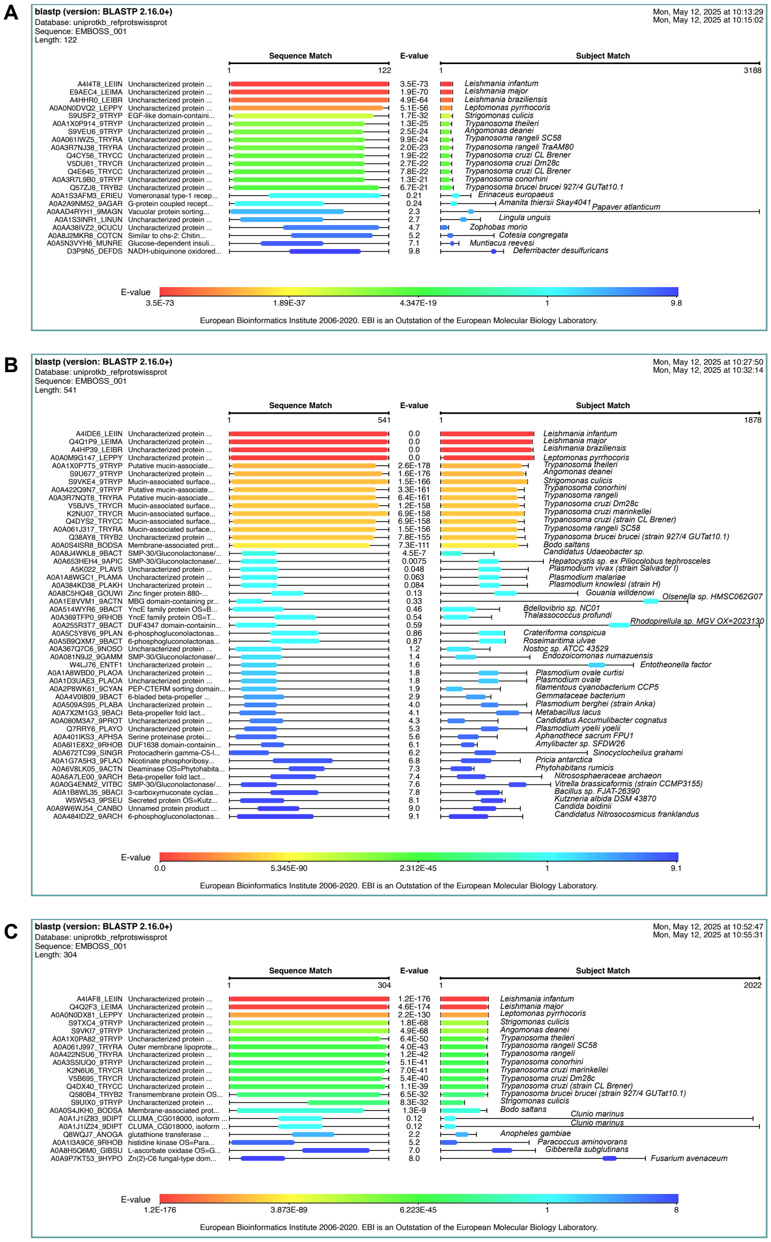


**Supplementary Figure 7.** BLASTP search in UniprotKB and SwissProt Reference proteins database for **(A)** LtVBP1, **(B)** LtVBP2, and **(C)** LtVBP3, showing protein conservation only in kinetoplastids.

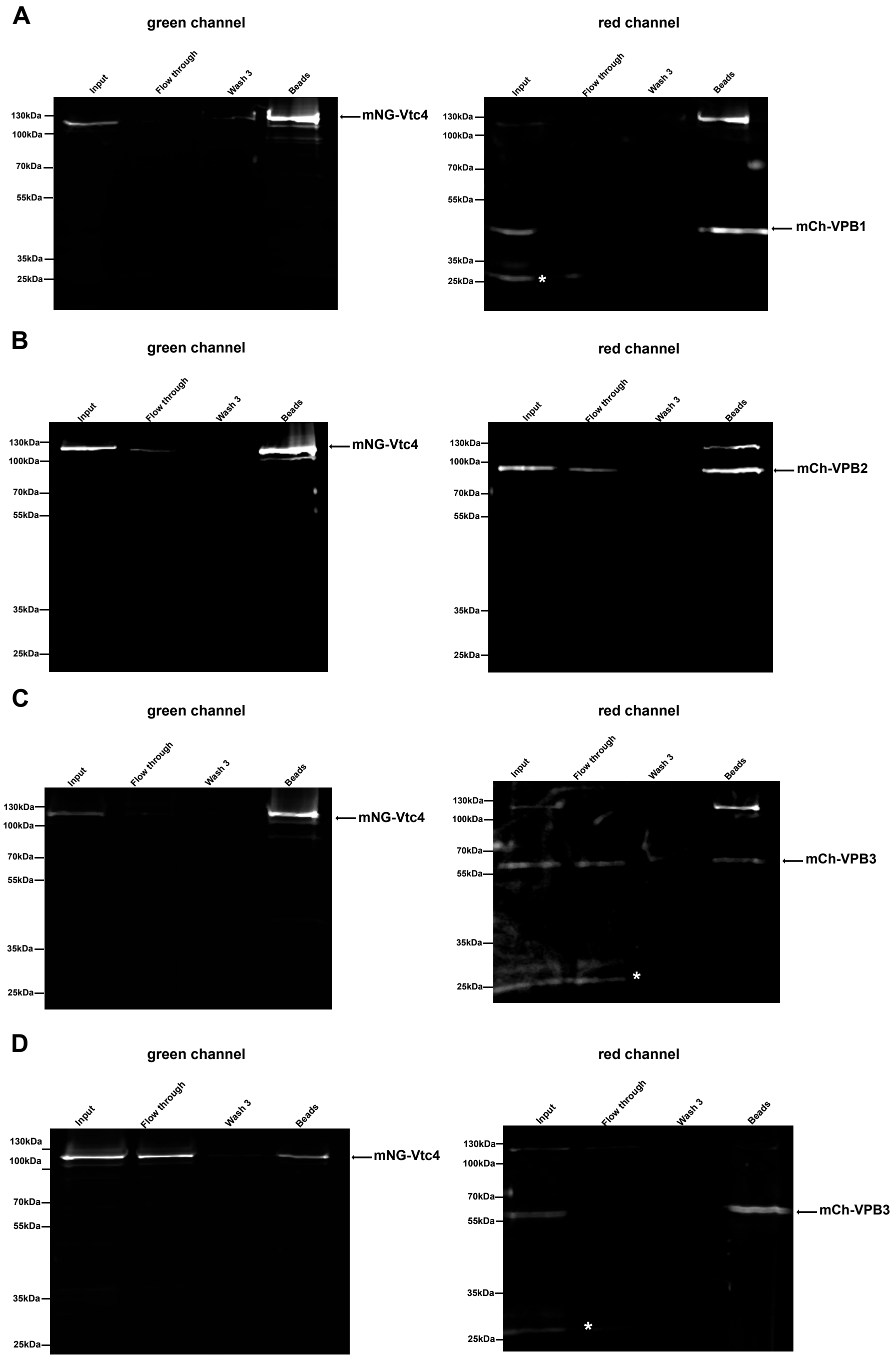


**Supplementary Figure 8.** Pulldown assays evaluating direct interactions between mNG-LtVtc4 and Novel VTC binding partners (VBPs1-3) tagged with mCh. **(A)** Detection of mNG-LtVtc4 signal in the green fluorescence channel (488 nm), indicating the presence of Vtc4 in the assay and detection of LtVBP1-mCh in the red fluorescence channel (532 nm) using mNG Trap Agarose Beads. **(B)** Detection of mNG-LtVtc4 signal in the green fluorescence channel (488 nm), indicating the presence of Vtc4 in the assay and detection of LtVBP2-mCh in the red fluorescence channel (532 nm) using mNG Trap Agarose Beads. **(C)** Detection of mNG-LtVtc4 signal in the green fluorescence channel (488 nm), indicating the presence of Vtc4 in the assay and detection of LtVBP3-mCh in the red fluorescence channel (532 nm) using mNG Trap Agarose Beads. **(D)** Detection of mNG-LtVtc4 signal in the green fluorescence channel (488 nm), indicating the presence of Vtc4 in the assay and detection of LtVBP3-mCh in the red fluorescence channel (532 nm) using RFP-Trap Magnetic Particles M270. The mNG signal may also be partially visible in red channel, depending on its relative intensity, which result in yellow bands reflecting overlapping fluorescence in the merged images. The fluorescence signal marked by a white asterisk in the gel corresponds to the mCh tag alone, visible only in the input due to partial degradation.

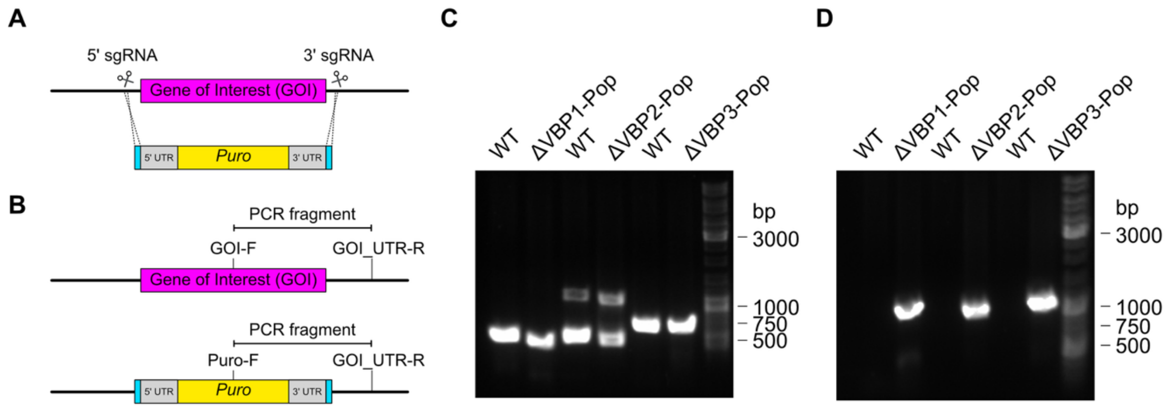


**S9 Fig. Strategy used for VBP knockout in *Leishmania tarentolae*.** **(A)** Schematic representation of the strategy used to knock out the VBP genes. Two sgRNAs, targeting the 5' and 3' ends of the coding sequence (CDS) of the gene of interest (GOI; scissors), were used to direct Cas9-mediated removal of the complete VBP CDS. A donor DNA containing a puromycin resistance marker (yellow) flanked by 30-nt homology arms (cyan) was also delivered to mediate double-strand break repair.

**(B)** Schematic representation of the two PCR assays used to screen for the presence or absence of the GOI and the puromycin resistance marker in the parental line and transfectant populations (ΔVBP1-pop, ΔVBP2-pop, and ΔVBP3-pop). **(C)** PCR amplification of fragments from the parental line and knockout transfectant populations using a GOI-specific forward primer (VBP1-F, VBP2-F, or VBP3-F) together with the corresponding 3'-UTR reverse primer (VBP1_UTR-R, VBP2_UTR-R, or VBP3_UTR-R). These primer pairs amplify fragments of 563, 568, and 739 bp, respectively. In all cases, both the parental line and knockout transfectant populations retained the gene. **(D)** PCR amplification of fragments from the parental line and knockout transfectant populations using the puromycin resistance marker forward primer (Puro-F) together with the corresponding 3'-UTR reverse primer (VBP1_UTR-R, VBP2_UTR-R, or VBP3_UTR-R). These primer pairs amplify fragments of 1204, 1200, and 1191 bp, respectively. In all cases, only the knockout transfectant populations produced the corresponding PCR fragments, whereas the parental line did not. Overall, the PCR amplification results shown in C and D indicate that the knockout transfectant populations contain the puromycin resistance cassette while still retaining the parental VBP gene signal. This suggests that these populations may contain cells with disrupted VBP loci together with cells or alleles retaining the parental VBP genes.

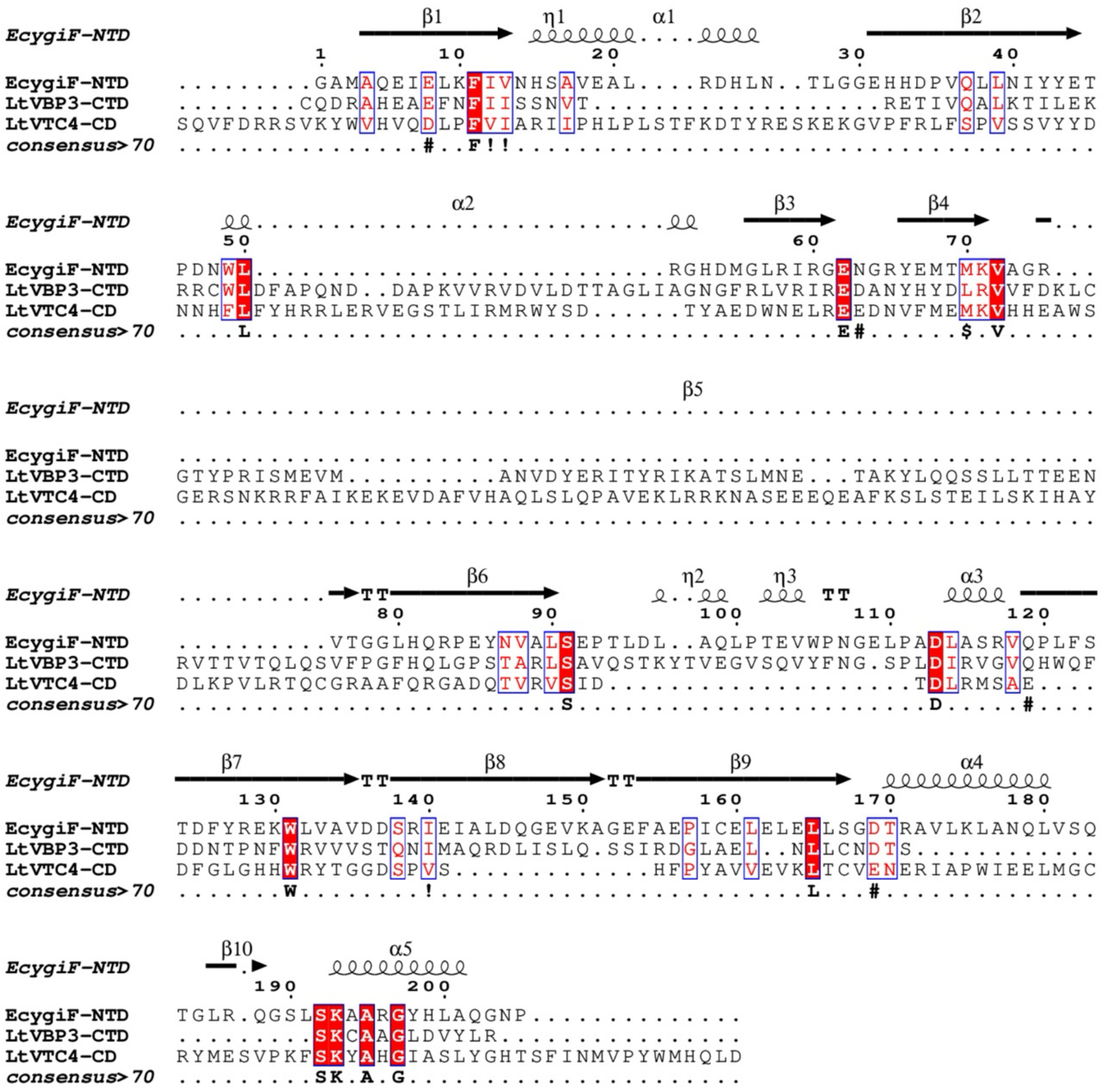

**Supplementary Figure 10.** Sequence alignment between LtVtc4-CD, LtVBP3-CTD, and EcygiF-NTD shows very low sequence identity: 10.6% between LtVtc4-CD and LtVBP3-CTD and 15.4% between LtVBP3-CTD and EcygiF-NTD. The red box shows sequence identity, and the blue rectangle shows consensus > 70%. The secondary structure corresponds to the solved structure of EcygiF [38].

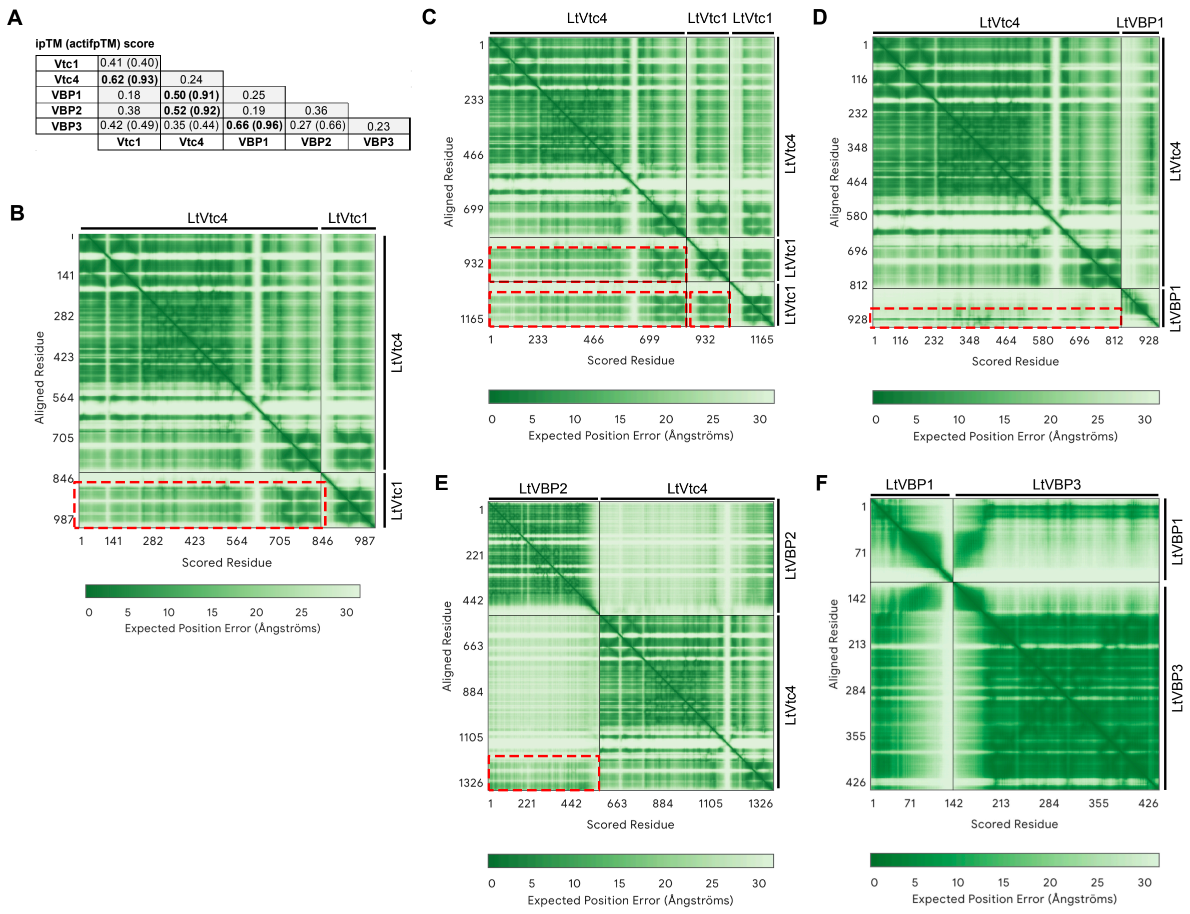


**S11 Fig. AlphaFold3 complex structure prediction of the VTC complex with its binding proteins. (A)** ipTM and activepTM scores for pairwise interactions among LtVtc1, LtVtc4, LtVBP1, LtVBP2, and LtVBP3. **(B-F)** PAE plot for complex structure prediction of: **(B)** LtVtc1 with LtVtc4, **(C)** two LtVtc1 with one LtVtc4, **(D)** LtVBP1 with LtVtc4, **(E)** LtVBP2 with LtVtc4, and **(F)** LtVBP1 with LtVBP3. The red-dashed rectangle highlights inter-subunit regions with low alignment error, indicating high confidence in interchain interactions.
