## Supplementary material for "Novel Binding Partners of the Vacuolar Transporter Chaperone (VTC) complex in Acidocalcisomes of *Leishmania tarentolae*": S6_Table

| **No.** | **Gene** | | **Protein name** | **MW (kDa)** | **Membrane association** | **Peptide counts (% Coverage)** | | | |
| --- | --- | --- | --- | --- | --- | --- | --- | --- | --- |
|  |  |  |  |  |  | **DDM** | | **FC12** | |
|  | ***L. tarentolae*** | ***T. brucei*** |  |  |  | **BirA-VTC1** | **BirA-VTC4** | **BirA-VTC1** | **BirA-VTC4** |
| 1 | LtaPh_0411200 | Tb927.9.8720 | Fructose-bisphosphatase | 39 | No | 4 (15) | 5 (23) | 4 (19) | 3 (12) |
| 2 | LtaPh_0503000 | Tb927.10.10610 | Protein tyrosine phosphatase, putative | 25 | Yes | 1 (5) | 3 (15) | 3 (15) | 2 (11) |
| 3 | LtaPh_0610500 | Tb927.7.5680 | 2-deoxy-D-ribose 5-phosphate aldolase | 28.4 | No | 1 (8) | 1 (8) | 0 (0) | 0 (0) |
| 4 | LtaPh_0800300 | Tb927.5.3560 | Vesicle-associated membrane protein, putative | 26.7 | No | 2 (15) | 0 (0) | 1 (6) | 1 (6) |
| 5 | LtaPh_1109700 | Tb927.11.6280 | Pyruvate phosphate dikinase | 100 | No | 19 (34) | 21 (35) | 12 (21) | 13 (23) |
| 6 | LtaPh_1302300 | Tb927.11.4550 | DUF4200 domain-containing protein (possible flagella associated protein) | 41.6 | No | 1 (6) | 1 (6) | 0 (0) | 0 (0) |
| 7 | LtaPh_1708500 | Tb927.5.2080, Tb927.10.16120 | GMP reductase | 66.7 | No | 5 (11) | 3 (10) | 1 (4) | 2 (8) |
| 8 | LtaPh_2102651 | Tb927.10.2020, Tb927.10.2010 | Phosphotransferase (hexokinase) | 51.7 | No | 21 (65) | 13 (42) | 12 (37) | 9 (30) |
| 9 | LtaPh_2109500 | Tb927.10.1390, Tb927.10.1400, Tb927.10.1470 | Hypoxanthine-guanine-xanthine phosphoribosyltransferase | 26.4 | No | 1 (8) | 1 (8) | 0 (0) | 0 (0) |
| 10 | LtaPh_2422400 | Tb927.8.6170 | Transketolase | 71.6 | No | 3 (8) | 1 (3) | 2 (5) | 1 (3) |
| 11 | LtaPh_2615900 | Tb927.2.4130 | Enoyl-CoA hydratase/Enoyl-CoA isomerase/3-hydroxyacyl-CoA dehydrogenase, putative | 88 | No | 3 (7) | 4 (6) | 2 (3) | 3 (7) |
| 12 | LtaPh_2704200 | Tb927.11.1020 | Ribokinase | 35.4 | No | 4 (19) | 3 (13) | 3 (13) | 2 (9) |
| 13 | LtaPh_2704700 | Tb927.11.1070 | Glycosomal transporter (GAT3), putative | 77.40 | Yes | 1 (3) | 0 (0) | 0 (0) | 0 (0) |
| 14 | LtaPh_2823400 | Tb927.11.11520 | Glycosomal membrane protein (PEX11) | 23.9 | Yes | 3 (20) | 5 (35) | 4 (23) | 4 (27) |
| 15 | LtaPh_2927100 | [Tb927.3.3270](https://tritrypdb.org/tritrypdb/app/record/gene/Tb927.3.3270) | ATP-dependent 6-phosphofructokinase | 54 | No | 9 (20) | 10 (24) | 7 (21) | 10 (27) |
| 16 | LtaPh_3001800 | Tb927.6.1500 | Alkyl dihydroxyacetonephosphate synthase | 70 | No | 12 (33) | 14 (35) | 4 (10 | 2 (5) |
| 17 | LtaPh_3105800 | Tb927.4.4070 | Mevalonate kinase | 35.7 | No | 1 (8) | 0 (0) | 0 (0) | 0 (0) |
| 18 | LtaPh_3328300 | Tb927.2.4130 | Enoyl-CoA hydratase/Enoyl-CoA isomerase/3-hydroxyacyl-CoA dehydrogenase, putative | 102 | No | 1 (1) | 1 (1) | 1 (1) | 0 (0) |
| 19 | LtaPh_3512600 | Tb927.5.930 | NADH-dependent fumarate reductas | 124 | No | 7 (10) | 6 (8) | 0 (0) | 0 (0) |
| 20 | LtaPh_3531000 | Tb927.9.12570 | Glycerol kinase, glycosomal | 56 | No | 2 (7) | 2 (4) | 3 (9) | 0 (0) |
| 21 | LtaPh_3537500 | Tb927.9.11600 | Gim5a protein, putative | 25 | No | 4 (31) | 3 (19) | 2 (16) | 2 (16) |
| 22 | LtaPh_3612400 | Tb927.10.5620 | Fructose-bisphosphate aldolase | 55 | No | 11 (34) | 6 (21) | 6 (21) | 6 (20) |
| 23 | LtaPh_3635700 | Tb927.11.9980, Tb927.11.1450 | 2-oxoglutarate dehydrogenase E1 component, putative | 114.7 | No | 6 (13) | 7 (14) | 9 (17) | 0 (0) |
| 24 | LtaPh_0600900 | Tb927.7.4770 | Cyclophilin-type peptidyl-prolyl cis-trans isomerase, putative | 23.7 | No | 0 (0) | 1 (10) | 2 (20) | 1 (10) |
| 25 | LtaPh_2607551 | Tb927.7.1130 | Trypanothione/Tryparedoxin dependent peroxidase 2 | 20.3 | No | 0 (0) | 1 (5) | 0 (0) | 0 (0) |
| 26 | LtaPh_3602500 | Tb927.9.12570 | Xylulokinase, putative | 54 | No | 0 (0) | 1 (5) | 0 (0) | 0 (0) |
| 27 | LtaPh_3631800 | Tb927.11.9420 | ATP synthase, putative | 25 | No | 0 (0) | 1 (10) | 0 (0) | 0 (0) |
| 28 | LtaPh_2718600 | Tb927.2.4210 | Phosphoenolpyruvate carboxykinase (ATP) | 58 | No | 22 (52) | 12 (29) | 14 (36) | 12 (32) |
