## Supplementary material for "Novel Binding Partners of the Vacuolar Transporter Chaperone (VTC) complex in Acidocalcisomes of *Leishmania tarentolae*": S7_Table

| **No.** | **Gene** | | **Protein name** | **MW (kDa)** | **Membrane association** | **Peptide counts (% Coverage)** | | | | |
| --- | --- | --- | --- | --- | --- | --- | --- | --- | --- | --- |
|  |  |  |  |  |  | **DDM** | | | **FC12** | |
|  | ***L. tarentolae*** | ***T. brucei*** |  |  |  | **BirA-VTC1** | **BirA-VTC4** | **BirA-VTC1** | | **BirA-VTC4** |
| 1 | LtaPh_0411400 | Tb927.9.8740 | Double RNA binding domain protein 3 | 35.4 | No | 2 (11) | 1 (7) | 0 (0) | | 0 (0) |
| 2 | LtaPh_1014000 | Tb927.8.4450 | RNA-binding protein, putative | 52.3 | No | 1 (3) | 2 (4) | 2 (5) | | 1 (3) |
| 3 | LtaPh_1802300 | Tb927.10.13720 | RNA-binding protein 29, putative | 43.5 | No | 0 (0) | 1 (4) | 0 (0) | | 0 (0) |
| 4 | LtaPh_2713900 | Tb927.11.1980 | Zinc finger protein family member, putative | 60 | No | 3 (6) | 3 (6) | 2 (4) | | 1 (2) |
| 5 | LtaPh_3312500 | Tb927.10.11760, Tb927.10.12660 | Pumilio/PUF RNA binding protein 6 | 91.8 | No | 1 (2) | 0 (0) | 0 (0) | | 0 (0) |
| 6 | LtaPh_3600300 | Tb927.10.4430 | Pumillo RNA binding protein PUF1 | 61 | No | 0 (0) | 0 (0) | 1 (3) | | 0 (0) |
| 7 | LtaPh_0512200 | Tb927.5.550 | V-type proton ATPase subunit D | 41.6 | No | 1 (8) | 0 (0) | 0 (0) | | 0 (0) |
| 8 | LtaPh_2300200 | Tb927.8.1960 | CCR4-NOT transcription complex subunit 11 (NO11, RNA degradation) | 48.7 | No | 3 (12) | 0 (0) | 1 (4) | | 0 (0) |
| 9 | LtaPh_2500900 | Tb927.9.10770 | Poly(A) binding protein, putative (PABP2) (Translation) | 61 | No | 8 (22) | 8 (21) | 4 (10) | | 11 (27) |
| 10 | LtaPh_2825100 | Tb927.11.11690 | Vacuolar proton pump subunit B | 55.6 | No | 2 (6) | 1 (3) | 1 (3) | | 0 (0) |
| 11 | LtaPh_3424900 | Tb927.4.2040, Tb927.4.2030 | Alba domain-containing protein (RNA binding) | 23 | No | 7 (43) | 6 (43) | 7 (43) | | 6 (43) |
| 12 | LtaPh_3425200 | Tb927.4.2000 | RuvB-like helicase | 54 | No | 1 (3) | 2 (7) | 0 (0) | | 0 (0) |
| 13 | LtaPh_3434000 | Tb927.4.1270 | RuvB-like helicase | 50 | No | 1 (4) | 3 (7) | 0 (0) | | 0 (0) |
| 14 | LtaPh_3435900 | Tb927.4.1080 | V-type ATPase, A subunit, putative | 67.7 | No | 1 (3) | 2 (5) | 2 (5) | | 1 (2) |
| 15 | LtaPh_3613500 | Tb927.10.5770 | Valosin-containing protein (AAA ATPase) | 86.9 | No | 4 (11) | 2 (5) | 0 (0) | | 0 (0) |
| 16 | LtaPh_3034700 | Tb927.6.4770 | Protein mkt1, putative | 91 | No | 0 (0) | 0 (0) | 1 (3) | | 0 (0) |
| 17 | LtaPh_3600300 | Tb927.10.4430 | Pumillo RNA binding protein PUF1 | 61 | No | 0 (0) | 0 (0) | 1 (3) | | 0 (0) |
