## Supplementary material for "Novel Binding Partners of the Vacuolar Transporter Chaperone (VTC) complex in Acidocalcisomes of *Leishmania tarentolae*": S8_Table

| **No.** | **Gene** | | **Protein name** | **MW (kDa)** | **Membrane association** | **Peptide counts (% Coverage)** | | | |
| --- | --- | --- | --- | --- | --- | --- | --- | --- | --- |
|  |  |  |  |  |  | **DDM** | | **FC12** | |
|  | ***L. tarentolae*** | ***T. brucei*** |  |  |  | **BirA-VTC1** | **BirA-VTC4** | **BirA-VTC1** | **BirA-VTC4** |
| 1 | LtaPh_0411200 | Tb927.9.8720 | Fructose-bisphosphatase | 39 | No | 4 (15) | 5 (23) | 4 (19) | 3 (12) |
| 2 | LtaPh_1109700 | Tb927.11.6280 | Pyruvate phosphate dikinase | 100 | No | 19 (34) | 21 (35) | 12 (21) | 13 (23) |
| 3 | LtaPh_2109500 | Tb927.10.1390, Tb927.10.1400, Tb927.10.1470 | Hypoxanthine-guanine-xanthine phosphoribosyltransferase | 26.4 | No | 1 (8) | 1 (8) | 0 (0) | 0 (0) |
| 4 | LtaPh_2927100 | [Tb927.3.3270](https://tritrypdb.org/tritrypdb/app/record/gene/Tb927.3.3270) | ATP-dependent 6-phosphofructokinase | 54 | No | 9 (20) | 10 (24) | 7 (21) | 10 (27) |
| 5 | LtaPh_3531000 | Tb927.9.12570 | Glycerol kinase, glycosomal | 56 | No | 2 (7) | 2 (4) | 3 (9) | 0 (0) |
| 6 | LtaPh_2718600 | Tb927.2.4210 | Phosphoenolpyruvate carboxykinase (ATP) | 58 | No | 22 (52) | 12 (29) | 14 (36) | 12 (32) |
| 7 | LtaPh_0605351 | Tb927.7.5180 | 60S ribosomal protein L23a, putative | 16.4 | No | 3 (30) | 5 (43) | 3 (30) | 1 (8) |
| 8 | LtaPh_0909431 | Tb927.11.13040, Tb927.11.13020,Tb927.11.13030 | Calmodulin | 16.80 | no | 7 (64) | 4 (34) | 6 (53) | 4 (39) |
| 9 | LtaPh_1012631 | Tb927.1.2430, Tb927.1.2450, Tb927.1.2470, Tb927.1.2490, Tb927.1.2510, Tb927.1.2530, Tb927.1.2550 | Histone domain-containing protein | 14.6 | No | 2 (18) | 1 (10) | 1 (10) | 1 (10) |
| 10 | LtaPh_1110600 | Tb927.11.6200, Tb927.11.6200 | 60S ribosomal protein L28, putative | 16.3 | No | 2 (11) | 3 (20) | 3 (20) | 3 (20) |
| 11 | LtaPh_1111000 | Tb927.11.6140, Tb927.3.4360 | 40S ribosomal protein S15a | 14.7 | No | 2 (13) | 2 (22) | 2 (13) | 2 (13) |
| 12 | LtaPh_1303351 | Tb927.11.4450 | Alba domain-containing protein | 13.3 | No | 3 (31) | 3 (31) | 3 (31) | 2 (26) |
| 13 | LtaPh_1304500 | Tb927.11.4290, Tb927.10.8430 | 40S ribosomal protein S12 | 24.8 | No | 2 (13) | 1 (8) | 2 (13) | 1 (8) |
| 14 | LtaPh_1413900 | Tb927.7.3630 | TPR-repeat-containing chaperone protein DNAJ, putative | 65 | No | 1 (3) | 0 (0) | 1 (2) | 0 (0) |
| 15 | LtaPh_1510200 | Tb927.10.11390 | 60S ribosomal protein L6, putative | 21.2 | No | 6 (34) | 5 (24) | 4 (29) | 4 (22) |
| 16 | LtaPh_1510481 | Tb927.9.5860, Tb927.9.5770 | Tryparedoxin peroxidase | 27.3 | No | 7 (29) | 4 (15) | 4 (16) | 4 (20) |
| 17 | LtaPh_1514200 | Tb927.9.5150 | Ribonucloprotein | 13.5 | No | 1 (14) | 1 (14) | 1 (19) | 1 (14) |
| 18 | LtaPh_1700771 | Tb927.10.2100, Tb927.10.2090, Tb927.10.2100 | Elongation factor 1-alpha | 61 | No | 15 (38) | 16 (35) | 0 (0) | 13 (33) |
| 19 | LtaPh_1802300 | Tb927.10.13720 | RNA-binding protein 29, putative | 43.5 | No | 0 (0) | 1 (4) | 0 (0) | 0 (0) |
| 20 | LtaPh_1806551 | Tb927.10.13500, Tb927.11.9710 | 60S ribosomal protein L10a, putative | 24.6 | No | 7 (43) | 0 (0) | 0 (0) | 0 (0) |
| 21 | LtaPh_1900201 | Tb927.10.10460, Tb927.10.10470, Tb927.10.10480, Tb927.10.10490, Tb927.10.10500, Tb927.10.10510, Tb927.10.10520, Tb927.10.10530, Tb927.10.10540, Tb927.10.10550, Tb927.10.10560, Tb927.10.10570, Tb927.10.10580, Tb927.10.10590 | Histone domain-containing protein | 12.3 | No | 6 (54) | 6 (54) | 5 (42) | 7 (54) |
| 22 | LtaPh_1900300 | Tb927.10.14710, Tb927.10.14600 | 40S ribosomal protein S2 | 28.2 | No | 6 (25) | 5 (25) | 0 (0) | 6 (33) |
| 23 | LtaPh_1900700 | Tb927.10.14750, Tb927.10.14630 | Fibrillarin, putative | 31.7 | No | 1 (7) | 1 (4) | 1 (4) | 1 (4) |
| 24 | LtaPh_2107401 | Tb927.10.1590, Tb927.9.15210 | 60S ribosomal protein L36, putative | 12 | No | 4 (37) | 4 (44) | 3 (37) | 3 (37) |
| 25 | LtaPh_2111061 | Tb927.7.2820, Tb927.7.2830, Tb927.7.2840, Tb927.7.2850, Tb927.7.2860, Tb927.7.2870, Tb927.7.2880, Tb927.7.2890, Tb927.7.2900, Tb927.7.2910, Tb927.7.2920, Tb927.7.2930, Tb927.7.2940 | Histone H2A | 14 | Yes | 4 (31) | 2 (19) | 4 (31) | 3 (26) |
| 26 | LtaPh_2204551 | Tb927.7.2370 | 40S ribosomal protein S15, putative | 20.1 | No | 3 (18) | 3 (15) | 1 (9) | 2 (11) |
| 27 | LtaPh_2215231 | Tb927.7.2370 | Ribosomal_L14e domain-containing protein | 17.5 | No | 2 (8) | 1 (5) | 2 (8) | 1 (5) |
| 28 | LtaPh_2300500 | Tb927.9.5860, Tb927.9.5770 | Tryparedoxin peroxidase | 25.5 | No | 1 (6) | 1 (6) | 1 (6) | 1 (6) |
| 29 | LtaPh_2422461 | Tb927.8.6150, Tb927.8.6160 | 40S ribosomal protein S8 | 25.2 | No | 2 (13) | 3 (19) | 2 (14) | 2 (14) |
| 30 | LtaPh_2509600 | Tb927.2.1560, Tb927.11.880 | Peptidyl-prolyl cis-trans isomerase | 28.1 | Yes | 1 (6) | 1 (6) | 2 (9) | 1 (6) |
| 31 | LtaPh_2525751 | Tb927.2.2670 | Histone H4 variant | 12.3 | No | 2 (11) | 2 (11) | 2 (11) | 2 (11) |
| 32 | LtaPh_2621200 | Tb927.9.2390, Tb927.9.2400 | Hypothetical protein, conserved | 23.2 | No | 1 (7) | 1 (12) | 1 (12) | 0 (0) |
| 33 | LtaPh_2623700 | Tb927.10.3280 | Ribosomal protein L38, putative | 9.5 | No | 1 (12) | 1 (12) | 1 (12) | 0 (0) |
| 34 | LtaPh_2802200 | Tb927.11.7350 | Histone domain-containing protein | 15.8 | No | 2 (18) | 0 (0) | 1 (8) | 0 (0) |
| 35 | LtaPh_2810300 | Tb927.11.7705 | 40S ribosomal protein S14 | 15.6 | No | 1 (13) | 0 (0) | 2 (24) | 2 (24) |
| 36 | LtaPh_2826361 | Tb927.11.11830 | 40S ribosomal protein S17, putative | 16.5 | No | 4 (29) | 2 (13) | 3 (22) | 1 (6) |
| 37 | LtaPh_2909000 | Tb927.3.3490 | High mobility group protein homolog tdp-1, putative | 34 | No | 5 (20) | 5 (20) | 6 (24) | 3 (15) |
| 38 | LtaPh_3029761 | Tb927.6.4300, Tb927.10.6880, Tb927.6.4280 | Glyceraldehyde 3-phosphate dehydrogenase, glycosomal | 39 | No | 11 (38) | 10 (35) | 11 (38) | 11 (37) |
| 39 | LtaPh_3204551 | Tb927.10.14580 | 60S ribosomal protein L17, putative | 19.1 | No | 3 (16) | 4 (23) | 1 (6) | 1 (6) |
| 40 | LtaPh_3317850 | Tb927.2.1560, Tb927.2.1680 | Peptidyl-prolyl cis-trans isomerase | 21.4 | No | 1 (5) | 1 (5) | 0 (0) | 1 (5) |
| 41 | LtaPh_3439500 | Tb927.4.750 | 50S ribosomal protein L7Ae, putative | 16.7 | No | 1 (18) | 0 (0) | 1 (6) | 1 (18) |
| 42 | LtaPh_3511300 | Tb927.5.800 | Casein kinase I, isoform 2 | 40 | No | 1 (3) | 1 (3) | 1 (3) | 1 (3) |
| 43 | LtaPh_3517200 | Tb927.8.6180, Tb927.9.14370 | 60S ribosomal protein L26, putative | 16 | No | 1 (5) | 1 (5) | 1 (7) | 2 (12) |
| 44 | LtaPh_3630021 | Tb927.10.7330, Tb927.10.7340 | 40S ribosomal protein S24e | 15.8 | No | 2 (20) | 3 (28) | 4 (45) | 3 (28) |
| 45 | LtaPh_3631451 | Tb927.10.7500 | Fibrillarin, putative | 31 | No | 2 (11) | 2 (12) | 1 (6) | 1 (6) |
| 46 | LtaPh_3635000 | Tb927.11.10025 | 60S ribosomal protein L29 | 18 | No | 2 (15) | 2 (15) | 2 (15) | 2 (15) |
| 47 | LtaPh_3638751 | Tb927.11.9710 | 60S ribosomal protein L10a | 29 | No | 0 (0) | 4 (21) | 3 (16) | 5 (26) |
| 48 | LtaPh_3650521 | Tb927.11.11010 | Hypothetical protein, conserved | 42.8 | No | 11 (37) | 11 (40) | 11 (39) | 9 (32) |
| 49 | LtaPh_3659900 | Tb927.10.8950 | Kinetoplast-associated protein 4 isoform 1 | 14.1 | No | 1 (10) | 1 (10) | 1 (10) | 1 (10) |
| 50 | LtaPh_3660700 | Tb927.10.8890 | Kinetoplast DNA-associated protein, putative | 18.5 | No | 2 (16) | 2 (16) | 1 (7) | 3 (29) |
