## Supplementary material for "Novel Binding Partners of the Vacuolar Transporter Chaperone (VTC) complex in Acidocalcisomes of *Leishmania tarentolae*": S9_Table

| **No.** | **Gene** | | **Protein name** | **MW (kDa)** | **Membrane association** | **Peptide counts (% Coverage)** | | | |
| --- | --- | --- | --- | --- | --- | --- | --- | --- | --- |
|  |  |  |  |  |  | **DDM** | | **FC12** | |
|  | ***L. tarentolae*** | ***T. brucei*** |  |  |  | **BirA-VTC1** | **BirA-VTC4** | **BirA-VTC1** | **BirA-VTC4** |
| 1 | LtaPh_0104700 | Tb927.9.4080 | Stumpy formation signalling pathway protein, putative | 177.6 | No | 1 (2) | 1 (2) | 1 (2) | 0 (0) |
| 2 | LtaPh_0107700 | Tb927.9.4820 | Calcium/potassium channel (CAKC), putative | 113.5 | Yes | 0 (0) | 1 (2) | 0 (0) | 0 (0) |
| 3 | LtaPh_0411200 | Tb927.9.8720 | Fructose-bisphosphatase | 39 | No | 4 (15) | 5 (23) | 4 (19) | 3 (12) |
| 4 | LtaPh_0503000 | Tb927.10.10610 | Protein tyrosine phosphatase, putative | 25 | Yes | 1 (5) | 3 (15) | 3 (15) | 2 (11) |
| 5 | LtaPh_0512200 | Tb927.5.550 | V-type proton ATPase subunit D | 41.6 | No | 1 (8) | 0 (0) | 0 (0) | 0 (0) |
| 6 | LtaPh_0600900 | Tb927.7.4770 | Cyclophilin-type peptidyl-prolyl cis-trans isomerase, putative | 23.7 | Yes | 0 (0) | 1 (10) | 2 (20) | 1 (10) |
| 7 | LtaPh_0602300 | Tb927.7.4900 | Exoribonuclease 2 | 169 | No | 0 (0) | 1 (1) | 1 (1) | 0 (0) |
| 8 | LtaPh_0606100 | Tb927.7.5230 | Lanosterol synthase, putative | 112.7 | Yes | 2 (3) | 1 (1) | 1 (1) | 1 (1) |
| 9 | LtaPh_0610500 | Tb927.7.5680 | 2-deoxy-D-ribose 5-phosphate aldolase | 28.4 | No | 1 (8) | 1 (8) | 0 (0) | 0 (0) |
| 10 | LtaPh_0800300 | Tb927.5.3560 | Vesicle-associated membrane protein, putative | 26.7 | No | 2 (15) | 0 (0) | 1 (6) | 1 (6) |
| 11 | LtaPh_0800800 | Tb927.5.3600 | ATP-dependent DEAD/H RNA helicase, putative | 108 | Yes | 1 (1) | 0 (0) | 0 (0) | 0 (0) |
| 12 | LtaPh_0902200 | Tb927.11.12220 | VTC4 | 93.1 | Yes | 25 (53.3) | 18 (30.3); 30 (35) | 5 (9.5) | 20 (32.4); 36 (39) |
| 13 | LtaPh_0902700 | Tb927.11.12270 | Drug resistence protein, putative | 66.5 | Yes | 0 (0) | 1 (2) | 0 (0) | 0 (0) |
| 14 | LtaPh_0903700 | Tb927.11.12360 | Ankyrin repeat protein, putative | 22.7 | No | 2 (11) | 1(11) | 0 (0) | 0 (0) |
| 15 | LtaPh_0904900 | Tb927.11.12490 | Hypothetical protein, conserved (possible potassium channel) | 79.5 | Yes | 0 (0) | 3 (7) | 0 (0) | 1 (2) |
| 16 | LtaPh_1002800 | Tb927.8.3690 | Isocitrate dehydrogenase [NADP] | 48.5 | Yes | 4(17) | 3(12) | 2(7) | 0 (0) |
| 17 | LtaPh_1015500 | Tb927.8.4610 | Ras-related protein RabX1 | 24.5 | No | 2(13) | 2(15) | 2 (13) | 0(0) |
| 18 | LtaPh_1108200 | Tb927.5.2750 | Alpha/beta hydrolase family, putative | 54.1 | No | 0(0) | 2(9) | 1(3) | 0(0) |
| 19 | LtaPh_1109700 | Tb927.11.6280 | Pyruvate phosphate dikinase | 100 | No | 19 (34) | 21 (35) | 12 (21) | 13 (23) |
| 20 | LtaPh_1302300 | Tb927.11.4550 | DUF4200 domain-containing protein (possible flagella associated protein) | 41.6 | No | 1 (6) | 1 (6) | 0 (0) | 0 (0) |
| 21 | LtaPh_1307700 | Tb927.11.3980 | Metallo-peptidase, Clan ME, Family M16 | 57.8 | No | 0 (0) | 2 (5) | 2 (7) | 0 (0) |
| 22 | LtaPh_1312500 | Tb927.11.3490 | Hypothetical protein, conserved (possible mRNA binding)) | 92 | No | 2 (4) | 1 (1) | 3 (5) | 0 (0) |
| 23 | LtaPh_1409100 | Tb927.7.3980 | Tc40 antigen-like (possible mRNA binding) | 91.8 | No | 2 (3) | 0 (0) | 2 (4) | 0 (0) |
| 24 | LtaPh_1410200 | Tb927.7.3900 | VTC1 | 20 | Yes | 5 (34.0); 21 (52) | 1 (8.3) | 2 (14.0); 17 (42) | 1 (8.3) |
| 25 | LtaPh_1512200 | Tb927.9.5490 | Mg^2+^ transporter, putative | 90.7 | Yes | 1 (2) | 2 (4) | 0 (0) | 0 (0) |
| 26 | LtaPh_1601400 | Tb927.5.3870 | LSU ribosomal protein, mitochondrial, putative | 36.5 | No | 0(0) | 1(6) | 1(6) | 0(0) |
| 27 | LtaPh_1706300 | Tb927.8.1160, Tb927.8.1180, Tb927.8.1200 | V-type Ca2+-ATPase, putative | 122.6 | Yes | 2 (3) | 3 (3) | 2 (3) | 1 (3) |
| 28 | LtaPh_1708500 | Tb927.5.2080, Tb927.10.16120 | GMP reductase | 66.7 | No | 5 (11) | 3 (10) | 1 (4) | 2 (8) |
| 29 | LtaPh_2011000 | Tb927.1.1930 | Phosphatidylinositol 3-kinase, putative | 360.4 | No | 1 (1) | 0 (0) | 0 (0) | 1 (1) |
| 30 | LtaPh_2102651 | Tb927.10.2020, Tb927.10.2010 | Phosphotransferase (hexokinase) | 51.7 | No | 21 (65) | 13 (42) | 12 (37) | 9 (30) |
| 31 | LtaPh_2108200 | Tb927.10.1510 | CCR4-NOT transcription complex subunit 1 | 249 | No | 1 (1) | 3 (2) | 1 (1) | 0 (0) |
| 32 | LtaPh_2109500 | Tb927.10.1390, Tb927.10.1400, Tb927.10.1470 | Hypoxanthine-guanine-xanthine phosphoribosyltransferase | 26.4 | No | 1 (8) | 1 (8) | 0 (0) | 0 (0) |
| 33 | LtaPh_2121161 | Tb927.10.200, Tb927.11.7480 | V-type proton ATPase subunit c | 19 | Yes | 1 (8) | 1 (8) | 1 (8) | 1 (8) |
| 34 | LtaPh_2211000 | Tb927.7.3180 | Adaptor complex AP-1 medium subunit, putative | 62.8 | No | 0 (0) | 0 (0) | 1 (2) | 0 (0) |
| 35 | LtaPh_2300200 | Tb927.8.1960 | CCR4-NOT transcription complex subunit 11 (NO11, RNA degradation) | 48.7 | No | 3 (12) | 0 (0) | 1 (4) | 0 (0) |
| 36 | LtaPh_2308300 | Tb927.8.2630 | Kinesin C | 86.3 | No | 7(13) | 9(14) | 2(3) | 4(9) |
| 37 | LtaPh_2319200 | Tb927.5.1300 | V-type proton ATPase subunit a | 87.8 | Yes | 2 (6) | 1 (3) | 1 (2) | 0 (0) |
| 38 | LtaPh_2422400 | Tb927.8.6170 | Transketolase | 71.6 | No | 3 (8) | 1 (3) | 2 (5) | 1 (3) |
| 39 | LtaPh_2500600 | Tb927.11.10980 | Vesicle transport protein | 17.2 | No | 1(14) | 0(0) | 1(14) | 0(0) |
| 40 | LtaPh_2500900 | Tb927.9.10770 | Poly(A) binding protein, putative (PABP2) (Translation) | 61 | No | 8 (22) | 8 (21) | 4 (10) | 11 (27) |
| 41 | LtaPh_2505700 | Tb927.11.550 | Scd6 (Suppressor of clathrin deficiency 6, putative) | 32.1 | No | 4 (21) | 3 (13) | 1 (5) | 1 (5) |
| 42 | LtaPh_2505900 | Tb927.11.570 | DnaJ domain containing protein, putative | 22 | No | 1(17) | 0(0) | 0(0) | 0(0) |
| 43 | LtaPh_2607551 | Tb927.7.1130 | Trypanothione/tryparedoxin dependent peroxidase 2 | 20.3 | No | 0 (0) | 1 (5) | 0 (0) | 0 (0) |
| 44 | LtaPh_2611300 | Tb927.7.740 | Chaperone protein DnaJ, putative | 71.4 | No | 1(2) | 0(0) | 0(0) | 0(0) |
| 45 | LtaPh_2615900 | Tb927.2.4130 | Enoyl-CoA hydratase/Enoyl-CoA isomerase/3-hydroxyacyl-CoA dehydrogenase, putative | 88 | No | 3 (7) | 4 (6) | 2 (3) | 3 (7) |
| 46 | LtaPh_2704200 | Tb927.11.1020 | Ribokinase | 35.4 | No | 4 (19) | 3 (13) | 3 (13) | 2 (9) |
| 47 | LtaPh_2704700 | Tb927.11.1070 | Glycosomal transporter (GAT3), putative | 77.40 | Yes | 1 (3) | 0 (0) | 0 (0) | 0 (0) |
| 48 | LtaPh_2713900 | Tb927.11.1980 | Zinc finger protein family member, putative | 60 | No | 3 (6) | 3 (6) | 2 (4) | 1 (2) |
| 49 | LtaPh_2717300 | Tb927.11.2320 | Hereditary spastic paraplegia protein strumpellin, putative | 143.5 | No | 0(0) | 1(4) | 0(0) | 0(0) |
| 50 | LtaPh_2718600 | Tb927.2.4210 | Phosphoenolpyruvate carboxykinase (ATP) | 58 | No | 22 (52) | 12 (29) | 14 (36) | 12 (32) |
| 51 | LtaPh_2810200 | Tb927.11.7740 | Dynein light chain, putative | 24.5 | No | 2 (18) | 1 (7) | 2 (12) | 0 (0) |
| 52 | LtaPh_2816900 | Tb927.11.9300 | Domain of unknown function (DUF3342), putative | 66.3 | No | 0(0) | 1(3) | 0(0) | 0(0) |
| 53 | LtaPh_2823400 | Tb927.11.11520 | Glycosomal membrane protein (PEX11) | 23.9 | Yes | 3 (20) | 5 (35) | 4 (23) | 4 (27) |
| 54 | LtaPh_2825100 | Tb927.11.11690 | Vacuolar proton pump subunit B | 55.6 | No | 2 (6) | 1 (3) | 1 (3) | 0 (0) |
| 55 | LtaPh_2829500 | Tb927.11.11160 | Pho91 | 81 | Yes | 1 (2) | 0 (0) | 0 (0) | 0 (0) |
| 56 | LtaPh_2923500 | Tb927.3.4720, Tb927.3.4760 | GTP-binding protein, putative; Dynamin-1-like | 77.2 | No | 7(15) | 8(16) | 5(10) | 3(6) |
| 57 | LtaPh_2927100 | [Tb927.3.3270](https://tritrypdb.org/tritrypdb/app/record/gene/Tb927.3.3270) | ATP-dependent 6-phosphofructokinase | 54 | No | 9 (20) | 10 (24) | 7 (21) | 10 (27) |
| 58 | LtaPh_2928300 | Tb927.3.3140 | Hypothetical protein, conserved | 13.6 | Yes | 1 (10) | 2 (25) | 2 (25) | 1 (10) |
| 59 | LtaPh_2930200 | Tb927.3.2950 | Ribonuclease inhibitor- like protein | 81.8 | No | 0(0) | 2(5) | 0(0) | 0(0) |
| 60 | LtaPh_2930400 | Tb927.3.2930 | RNA-binding protein RBP6, putative | 23.10 | No | 0 (0) | 0 (0) | 0 (0) | 1 (10) |
| 61 | LtaPh_3001800 | Tb927.6.1500 | Alkyl dihydroxyacetonephosphate synthase | 70 | No | 12 (33) | 14 (35) | 4 (10 | 2 (5) |
| 62 | LtaPh_3003900 | Tb927.6.1750 | Rab-GAP TBC domain-containing protein | 92.8 | No | 0(0) | 1(4) | 0(0) | 0(0) |
| 63 | LtaPh_3014400 | Tb927.6.2810 | ABC transporter, putative | 58.1 | No | 0(0) | 1(5) | 0(0) | 0(0) |
| 64 | LtaPh_3029761 | Tb927.6.4300, Tb927.10.6880, Tb927.6.4280 | Glyceraldehyde 3-phosphate dehydrogenase, glycosomal | 39 | No | 11 (38) | 10 (35) | 11 (38) | 11 (37) |
| 65 | LtaPh_3034700 | Tb927.6.4770 | Protein mkt1, putative | 91 | No | 0 (0) | 0 (0) | 1 (3) | 0 (0) |
| 66 | LtaPh_3100200 | Tb927.10.14160, Tb927.10.14170, Tb927.6.1520 | Aquaporin 9, putative | 34.7 | Yes | 0 (0) | 0 (0) | 1 (3) | 0 (0) |
| 67 | LtaPh_3105800 | Tb927.4.4070 | Mevalonate kinase | 35.7 | No | 1 (8) | 0 (0) | 0 (0) | 0 (0) |
| 68 | LtaPh_3111800 | Tb927.4.4380 | mPPase | 83.6 | Yes | 4 (6) | 4 (7) | 5 (7) | 5 (6) |
| 69 | LtaPh_3112621 | Tb927.4.4490 | Multidrug resistance protein E | 148.4 | No | 0(0) | 1(1) | 0(0) | 0(0) |
| 70 | LtaPh_3128400 | Tb927.4.4960, Tb927.8.7460 | ZnT | 47.2 | Yes | 3 (17) | 2 (8) | 1 (3) | 1 (3) |
| 71 | LtaPh_3134500 | Tb927.8.7110 | Serine/threonine-protein kinase NEK12.1, putative | 49.8 | No | 0(0) | 1(4) | 0(0) | 0(0) |
| 72 | LtaPh_3135300 | Tb927.11.16970, Tb927.11.11940 | Hypothetical protein, conserved | 27.5 | No | 0(0) | 2(20) | 0(0) | 0(0) |
| 73 | LtaPh_3207100 | Tb927.11.13900 | Centrin-5 | 22.8 | No | 1 (11) | 0 (0) | 0 (0) | 0 (0) |
| 74 | LtaPh_3302521 | Tb927.10.10800 | Palmitoyl acyltransferase 2, putative | 58.2 | Yes | 2 (6) | 1 (3) | 1 (3) | 1 (3) |
| 75 | LtaPh_3328300 | Tb927.2.4130 | Enoyl-CoA hydratase/Enoyl-CoA isomerase/3-hydroxyacyl-CoA dehydrogenase, putative | 102 | No | 1 (1) | 1 (1) | 1 (1) | 0 (0) |
| 76 | LtaPh_3330400 | Tb927.2.3610 | Mitochondrial ATP synthase subunit, putative | 17.2 | Yes | 1(13) | 1(13) | 1(13) | 0(0) |
| 77 | LtaPh_3405200 | Tb927.10.2880 | Voltage-dependent calcium channel subunit, putative | 312.2 | No | 0(0) | 1(1) | 0(0) | 0(0) |
| 78 | LtaPh_3410000 | Tb927.4.3540 | Hypothetical protein | 59 | No | 0 (0) | 1 (4) | 0 (0) | 0 (0) |
| 79 | LtaPh_3424900 | Tb927.4.2040, Tb927.4.2030 | Alba domain-containing protein (RNA binding) | 23 | No | 7 (43) | 6 (43) | 7 (43) | 6 (43) |
| 80 | LtaPh_3425200 | Tb927.4.2000 | RuvB-like helicase | 54 | No | 1 (3) | 2 (7) | 0 (0) | 0 (0) |
| 81 | LtaPh_3434000 | Tb927.4.1270 | RuvB-like helicase | 50 | No | 1 (4) | 3 (7) | 0 (0) | 0 (0) |
| 82 | LtaPh_3435900 | Tb927.4.1080 | V-type ATPase, A subunit, putative | 67.7 | No | 1 (3) | 2 (5) | 2 (5) | 1 (2) |
| 83 | LtaPh_3438300 | Tb927.4.860 | Hypothetical protein, conserve | 34.4 | Yes | 3 (13) | 0 (0) | 1 (6) | 1 (6) |
| 84 | LtaPh_3443000 | Tb927.4.410 | Cell differentiation protein-like protein | 54.1 | No | 0 (0) | 0 (0) | 1 (3) | 1 (3) |
| 85 | LtaPh_3505600 | Tb927.10.3990 | ATP-dependent DEAD-box RNA helicase, putative (DHH1) | 46.4 | No | 8 (29) | 6 (19) | 4 (13) | 4 (18) |
| 86 | LtaPh_3512600 | Tb927.5.930 | NADH-dependent fumarate reductas | 124 | No | 7 (10) | 6 (8) | 0 (0) | 0 (0) |
| 87 | LtaPh_3522500 | Tb927.9.13990 | RNA-binding protein, putative | 30 | No | 7 (36) | 8 (37) | 6 (36) | 9 (47) |
| 88 | LtaPh_3531000 | Tb927.9.12570 | Glycerol kinase, glycosomal | 56 | No | 2 (7) | 2 (4) | 3 (9) | 0 (0) |
| 89 | LtaPh_3537500 | Tb927.9.11600 | Gim5a protein, putative | 25 | No | 4 (31) | 3 (19) | 2 (16) | 2 (16) |
| 90 | LtaPh_3548100 | Tb927.9.9710 | Histidine phosphatase superfamily (branch 1), putative | 36.7 | Yes | 0 (0) | 1 (5) | 2 (9) | 1 (5) |
| 91 | LtaPh_3600300 | Tb927.10.4430 | Pumillo RNA binding protein PUF1 | 61 | No | 0 (0) | 0 (0) | 1 (3) | 0 (0) |
| 92 | LtaPh_3602500 | Tb927.9.12570 | Xylulokinase, putative | 54 | No | 0 (0) | 1 (5) | 0 (0) | 0 (0) |
| 93 | LtaPh_3612400 | Tb927.10.5620 | Fructose-bisphosphate aldolase | 55 | No | 11 (34) | 6 (21) | 6 (21) | 6 (20) |
| 94 | LtaPh_3613500 | Tb927.10.5770 | Valosin-containing protein (AAA ATPase) | 86.9 | No | 4 (11) | 2 (5) | 0 (0) | 0 (0) |
| 95 | LtaPh_3617400 | Tb927.10.6180 | FLA1-like protein | 60 | Yes | 8 (25) | 5 (14) | 5 (16) | 3 (9) |
| 96 | LtaPh_3621200 | Tb927.10.6610 | Chaperone protein DnaJ, putative | 33.9 | No | 0(0) | 2(9) | 0(0) | 0(0) |
| 97 | LtaPh_3625400 | Tb927.10.7020 | Acid phosphatase, putative | 54.3 | No | 0 (0) | 1 (5) | 0 (0) | 0 (0) |
| 98 | LtaPh_3631800 | Tb927.11.9420 | ATP synthase, putative | 25 | No | 0 (0) | 1 (10) | 0 (0) | 0 (0) |
| 99 | LtaPh_3631800 | Tb927.11.9420 | ATP synthase, putative | 25.3 | Yes | 0(0) | 1(10) | 0(0) | 0(0) |
| 100 | LtaPh_3635700 | Tb927.11.9980, Tb927.11.1450 | 2-oxoglutarate dehydrogenase E1 component, putative | 114.7 | No | 6 (13) | 7 (14) | 9 (17) | 0 (0) |
| 101 | LtaPh_3639600 | Tb927.11.9630 | Snf7, putative | 25.5 | No | 1(4) | 1(4) | 1(4) | 3(15) |
| 102 | LtaPh_3650521 | Tb927.11.11010 | Hypothetical protein, conserved | 42.8 | No | 11 (37) | 11 (40) | 11 (39) | 9 (32) |
| 103 | LtaPh_3654000 | Tb927.11.10630 | Hypothetical protein, conserved (possibly protein phosphatase regulator activity) | 114 | No | 1 (1) | 0 (0) | 0 (0) | 1 (1) |
